# CoTRA: a comprehensive R/Shiny framework for transparent bulk and single-cell RNA-seq analysis

**DOI:** 10.64898/2026.08.25.747017

**Authors:** Umair Seemab, Katri Vainionpaa, Ziaurrehman Tanoli, Henri Leinonen

## Abstract

Bulk RNA-seq and single-cell RNA-seq (scRNA-seq) are widely used to investigate gene-expression changes, but downstream analysis often requires multiple statistical, visualization, and reporting tools, creating fragmented workflows that are difficult to configure and reproduce. We developed CoTRA (Comprehensive Toolbox for RNA-seq Analysis), an open-source R/Shiny package providing independent bulk and scRNA-seq workflows within a common graphical environment. CoTRA supports quality control, differential expression, annotation, enrichment, dimensionality reduction, clustering, marker detection, cell-type annotation, differential abundance, trajectory inference, pathway activity, cell-cell communication, and reporting while exposing key analytical parameters. Compared with 14 other platforms across 49 predefined criteria, CoTRA fully supported 46 and partially supported three. Under matched inputs and parameters, CoTRA reproduced direct DESeq2, edgeR, and Seurat implementations, including identical significant bulk gene sets and scRNA-seq clustering (ARI = 1.000; NMI = 1.000). Retinal case studies recapitulated degeneration-associated transcriptional changes and demonstrated cell-type-resolved analysis. Synthetic scRNA-seq benchmarking scaled to 50,000 cells with 5.10 GB peak memory. CoTRA v1.0.0 requires R ≥ 4.4.0, has been tested on Linux, Windows, and macOS, is GPL-3 licensed, and is available at https://github.com/UmairSeemab/CoTRA, and support is provided through GitHub Issues.

**AUTHOR SUMMARY:** Modern sequencing technologies can measure the activity of thousands of genes across whole tissues or individual cells, but analyzing these data often requires researchers to combine many separate software tools and write substantial amounts of code. We developed CoTRA to make this process more accessible while keeping important analytical choices visible to the user. CoTRA provides graphical workflows for both bulk and single-cell RNA sequencing, covering data quality assessment, identification of expression changes, biological interpretation, cell clustering and annotation, and several advanced single-cell analyses. We tested CoTRA using published retinal datasets, compared its outputs with direct scripted analyses, and measured its computational performance as dataset size increased. The graphical workflows reproduced the corresponding scripted results when the same data and parameters were used, and the retinal examples recovered expected disease-associated expression patterns. CoTRA runs locally, so researchers do not need to upload their expression data to a mandatory external service. We hope that this combination of accessibility, transparency, and reproducible outputs will make transcriptomic analysis easier to use and inspect across collaborative biomedical research projects.

## INTRODUCTION

RNA sequencing (RNA-seq) and single-cell RNA sequencing (scRNA-seq) are widely used to investigate gene-expression changes across tissues, experimental conditions, and individual cell populations[1,2]. Their downstream analysis, however, often requires researchers to combine multiple statistical packages, visualization tools, data structures, and reporting procedures within a single project [3,4]. Bulk RNA-seq workflows may involve quality assessment, normalization, differential expression, annotation, enrichment, and visualization, whereas scRNA-seq analysis additionally requires cell-level quality control, dimensionality reduction, clustering, marker identification, cell-type annotation, and increasingly specialized analyses such as trajectory inference, pathway activity, differential abundance, and cell-cell communication [5–17].

Established methods are available for most of these tasks, but combining them commonly requires programming experience, management of package dependencies and intermediate data formats, and careful tracking of parameters across multiple analysis steps. These requirements can limit data reuse by researchers without continuous bioinformatics support and can lead to workflows fragmented across scripts, spreadsheets, standalone applications, and web services [18–26]. Such fragmentation also makes it more difficult to preserve intermediate outputs, document analytical decisions, and reproduce an analysis using the same computational settings.

Several open-source platforms have lowered the technical barrier to RNA-seq analysis, but they differ substantially in scope and workflow coverage. General or bulk-oriented environments include Galaxy [27], iDEP [28], BioJupies [29], DEBrowser [30], ideal [31], Glimma [32], and DiVenn [33], whereas Seurat [34], iSEE [35], ShiVA [36], and SinglePointRNA [37] focus primarily on single-cell analysis. BingleSeq [38], SEQUIN [39], and IRIS-EDA [40] provide both bulk and single-cell RNA-seq functionality, but with less comprehensive coverage of the combined analytical, downstream, export, and reproducibility features assessed in our comparison. Across these platforms, support varies in deployment, input formats, parameter configurability, visualization, functional interpretation, advanced single-cell analyses, reporting, and reproducibility-oriented outputs. This motivated the development of CoTRA as a locally deployable R/Shiny environment that supports independent bulk and scRNA-seq workflows within a common interface while retaining visible analytical choices, alternative established methods, and reusable figures, tables, processed objects, and session information.

To address this need, we developed CoTRA (Comprehensive Toolbox for RNA-seq Analysis), an open-source R package with a Shiny-based graphical interface for downstream bulk and single-cell RNA-seq analysis. CoTRA does not introduce a new statistical algorithm. Instead, it organizes established bioinformatics methods within modular workflows while exposing key analytical parameters and providing alternative methods at selected stages. The software can be executed on personal workstations or compatible remote computing infrastructure without mandatory submission of expression data to an external server. In this study, we describe the design and implementation of CoTRA and evaluate its functional scope relative to existing transcriptomic analysis platforms, its quantitative agreement with corresponding scripted implementations, its computational performance, and its application to representative bulk and single-cell RNA-seq datasets.

## DESIGN AND IMPLEMENTATION

### CoTRA architecture and workflow

CoTRA (Comprehensive Toolbox for RNA-seq Analysis) was developed as an installable R package with a Shiny-based graphical interface for downstream bulk and single-cell RNA-seq analysis. Bulk and single-cell RNA-seq are implemented as independent analytical branches within the same application. Analyses can be executed locally or in compatible remote R environments, including cloud-based Linux virtual machines running RStudio Server, without mandatory transfer of expression data to an external CoTRA-hosted server.

The software uses modular architecture comprising reusable R functions, Shiny user-interface and server components, dependency checks, analysis modules, reporting functions, and export utilities. CoTRA is launched through `runCoTRA()`, which creates a temporary writable application directory and directs generated outputs to a user-defined location or, by default, to a CoTRA_Results directory in the user’s home directory. CoTRA has been tested on Linux, Windows, and macOS. The current workflow begins with processed expression data and does not perform FASTQ preprocessing or read alignment. The overall architecture, supported inputs, analytical stages, and outputs are summarized in **Fig 1**.

**Fig 1.**
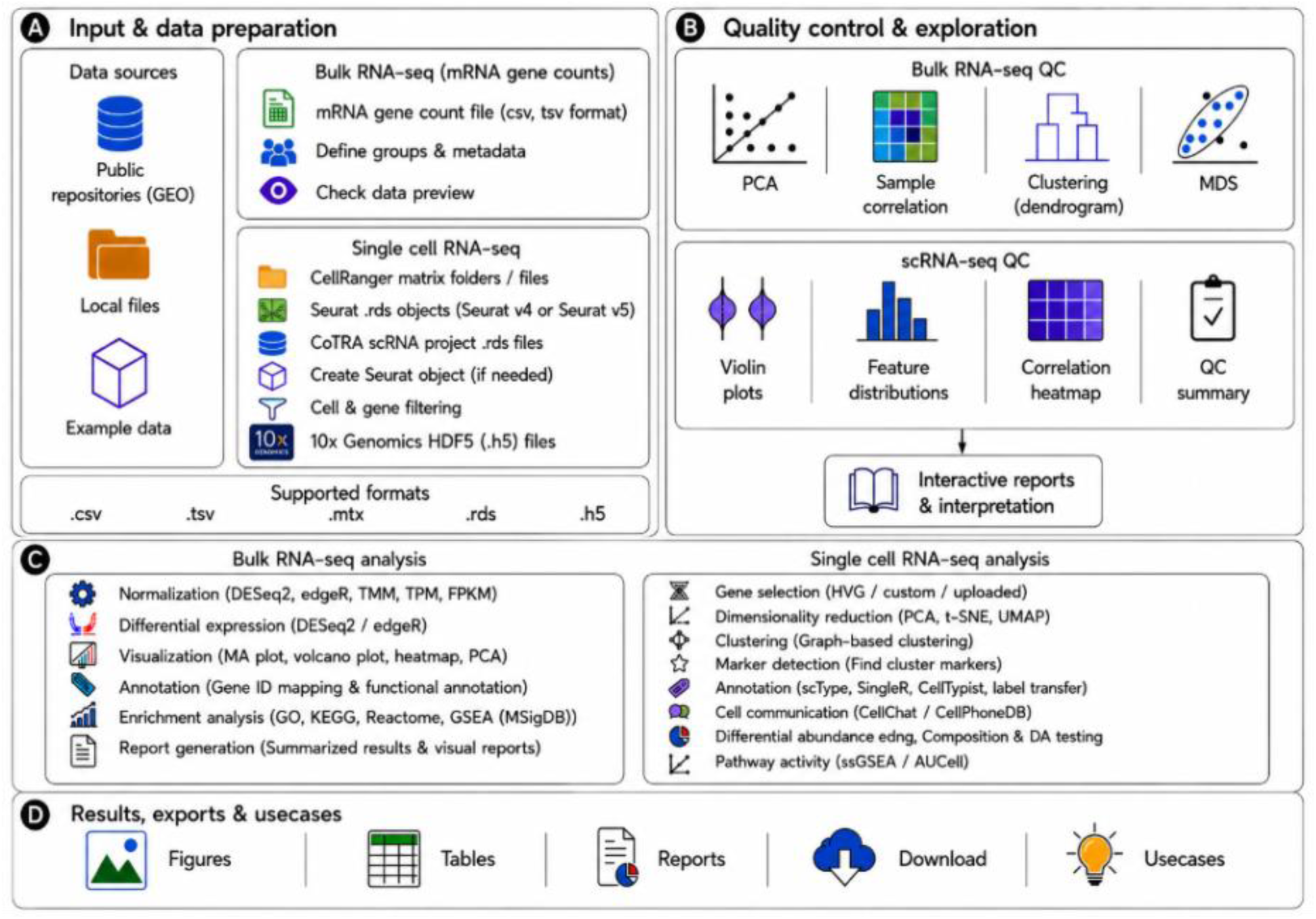
Overview of the CoTRA workflow for bulk and single-cell RNA-seq analysis. **(A)** CoTRA accepts bulk count matrices and single-cell inputs including Cell Ranger outputs, 10x HDF5 files, Seurat RDS objects, and CoTRA project files. **(B)** Quality control and exploratory modules provide PCA, correlation, clustering, MDS, and cell-level QC visualizations. **(C)** Dedicated bulk and scRNA-seq workflows support differential expression, annotation, enrichment, clustering, marker analysis, trajectory, pathway activity, differential abundance, and cell–cell communication. **(D)** Results can be exported as figures, tables, reports, gene lists, processed objects, and session files for reproducible downstream analysis.

### Bulk RNA-seq workflow

The bulk workflow accepts genes by sample count matrices in CSV, TSV, TXT, or TAB format. CoTRA detects delimiters and candidate gene-identifier columns, removes empty columns, converts expression columns to numeric values, and allows users to rename samples and define experimental groups. Raw counts are retained for differential expression testing, whereas TPM, FPKM, RPKM, and TMM normalized values are available for exploratory analysis and export when the required information is available. Differential expression can be performed with DESeq2 or edgeR. Downstream functions include sample-level quality assessment, PCA, multidimensional scaling, correlation analysis, hierarchical clustering, genomic annotation, and functional enrichment using Gene Ontology, KEGG, Reactome, and MSigDB Hallmark gene sets. Results are returned as visualizations together with downloadable tables and gene lists. Detailed defaults and configurable parameters are provided in **S1 Table**.

### Single-cell RNA-seq workflow

The single-cell workflow accepts Cell Ranger matrix directories, 10x Genomics HDF5 files, Seurat RDS objects, and CoTRA project RDS files. Multiple HDF5 files can be imported in a single analysis, with each file treated as a biological sample. CoTRA allows sample identifiers and experimental conditions to be assigned through the interface, prefixes cell barcodes with sample-specific identifiers to preserve uniqueness and combines the matrices before construction of the joint Seurat object. Imported data are converted to or validated as Seurat objects, with compatibility handling for Seurat v4 assay slots and Seurat v5 assay layers. Quality control includes detected features, total RNA counts, mitochondrial transcript percentage, and ribosomal transcript percentage, with user-adjustable thresholds. Preprocessing supports Seurat LogNormalize and SCTransform, followed by highly variable gene selection, PCA, UMAP or t-SNE, graph construction, and clustering. Marker analysis supports Wilcoxon, MAST, logistic regression, and DESeq2. Cell-type annotation can be performed using SingleR, custom marker-based scoring, imported CellTypist predictions, or Seurat label transfer. Advanced modules implement trajectory inference with Monocle3 or Slingshot, pathway activity with UCell, AUCell, GSVA, or VISION, cell-cell communication with CellChat, and differential-abundance approaches including replicate-aware and contingency-table-based methods. Detailed method settings are provided in **S1 Table**.

### Parameter transparency, reproducibility, and reporting

CoTRA keeps major analytical decisions visible through module-specific controls, including quality-control thresholds, differential-expression cutoffs, principal-component selection, clustering resolution, method selection, and downstream settings. Tables can be exported as CSV or XLSX files and figures as PNG, PDF, or SVG. Selected modules additionally generate reports, ZIP archives, processed RDS or Seurat objects, CoTRA project/session files, and session information. Fixed random seeds are used for selected stochastic single-cell procedures where supported. These outputs are intended to preserve analytical results and computational information for subsequent inspection and re-analysis. Interface examples and Help and Interpretation components are shown in **S1 Fig**.

### Performance and implementation evaluation

Computational performance was evaluated using synthetic scaling datasets and real retinal bulk and single-cell RNA-seq datasets. Each condition was executed in five independent R processes under single-thread constraints, with analytical runtime measured around the computational backend and peak resident memory obtained using GNU /usr/bin/time -v; Shiny user interaction and browser rendering were not included. Implementation reproducibility was assessed by comparing CoTRA outputs with direct scripted DESeq2, edgeR, and Seurat analyses using identical input data, parameters, filtering procedures, and contrasts. Concordance was evaluated using numerical agreement, gene-set overlap, correlation of effect-size estimates, and, for scRNA-seq, highly variable gene overlap, PCA concordance, Adjusted Rand Index (ARI), Normalized Mutual Information (NMI), and marker-set agreement. Detailed analytical settings are provided in **S1 Table**; the computational environment and benchmark results in **S2-S7 Tables**; implementation-concordance results in **S8-S9 Tables**; and test-data settings and expected outputs in **S10 Table**. Run-level and processed benchmarking resources are provided in **S1–S3 Data** and **S1 Code**, while implementation validation outputs and scripts are provided in **S4 Data** and **S2 Code**.

### Software comparison design

CoTRA was compared with 14 representative transcriptomic analysis platforms using 49 predefined criteria spanning deployment and usability, input handling, bulk RNA-seq, single-cell RNA-seq, visualization, biological interpretation, advanced analysis, and export/reproducibility. Evidence was obtained from direct software evaluation, official documentation, and published descriptions. Two evaluators independently assessed each criterion, with disagreements resolved by consensus before consolidation. Each criterion was classified as fully available, partially available, absent, or not applicable according to predefined definitions. The complete criterion-level comparison, evidence sources, software versions, and evaluation dates are provided in **S11 Table**, while the feature-comparison heatmap is presented in **S2 Fig**.

## RESULTS

### Bulk RNA-seq case study

To evaluate the bulk RNA-seq workflow, CoTRA was applied to a previously published retinal transcriptomic dataset comparing retinas from the rd10 mouse model of retinal degeneration (n = 8) with sex-matched wild-type (WT) retinas (n = 4) [41]. The dataset is available under GEO accession GSE238217 as part of SuperSeries GSE238218. The gene-count matrix was analyzed through sample level quality assessment, differential-expression analysis, genomic annotation, expression visualization, and functional enrichment (**Fig 2**). The corresponding count matrix is distributed with the CoTRA repository as a test dataset, and the parameters used to reproduce the analysis are provided in the Supplementary Information.

**Fig 2.**
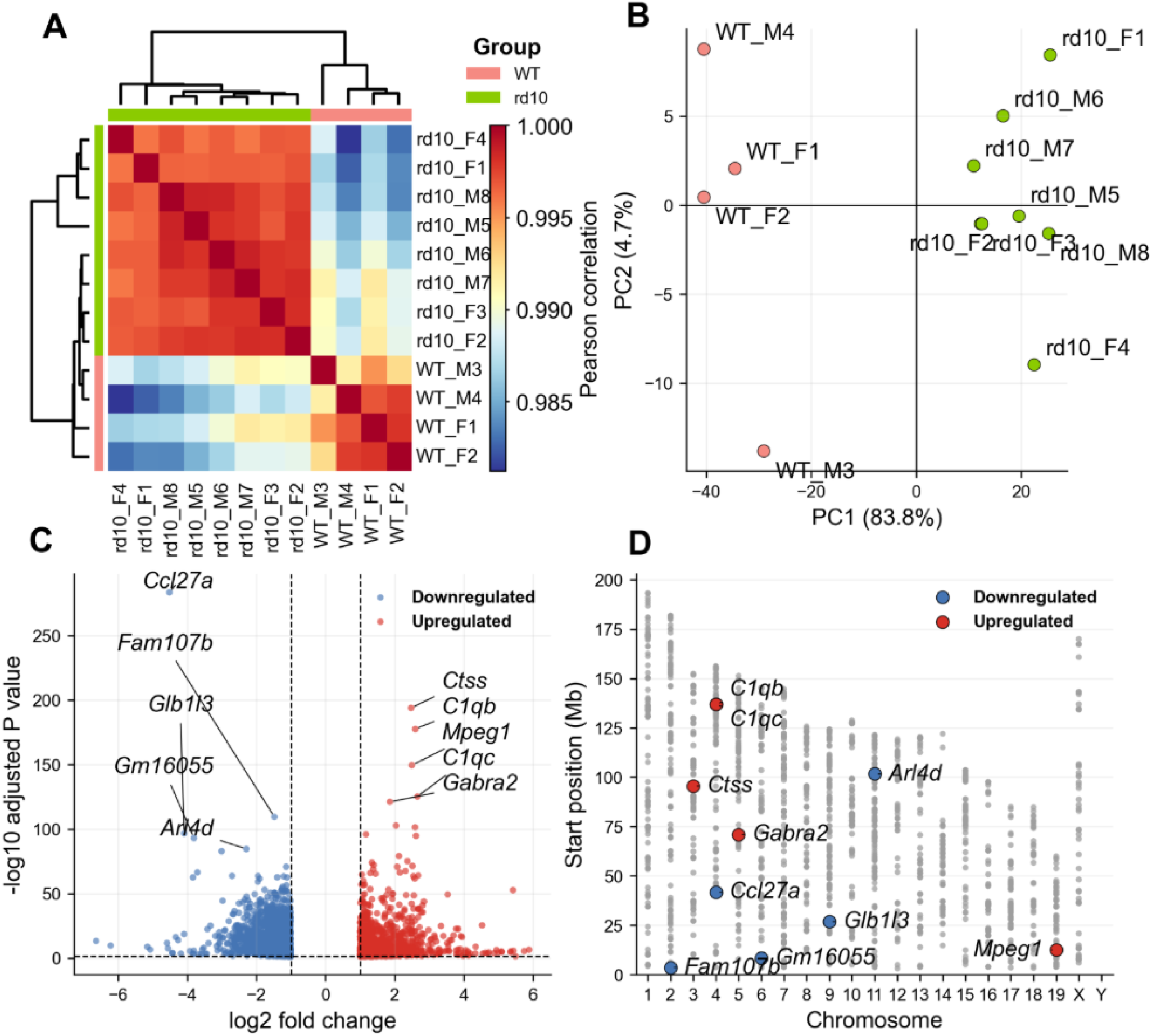
CoTRA-based bulk RNA-seq analysis of the rd10 retinal degeneration model. **(A)** Sample correlation heatmap showing strong within-group similarity and separation of rd10 (n = 8) and WT (n = 4) retinas. **(B)** PCA of normalized expression data by experimental group. **(C)** Volcano plot of rd10 versus WT differential expression using adjusted P < 0.05 and |log2 fold change| ≥ 1. **(D)** Chromosomal distribution of significant DEGs across mouse chromosomes, with selected upregulated and downregulated genes highlighted.

Sample-level analysis showed high within-group similarity and clear separation of rd10 and WT samples in the correlation heatmap (**Fig 2A**). PCA independently separated the two groups, with PC1 and PC2 explaining 83.8% and 4.7% of the variance, respectively (**Fig 2B**).

Using DESeq2 with adjusted P < 0.05 and |log2 fold change| ≥ 1, CoTRA identified 2,518 differentially expressed genes (DEGs), comprising 1,143 upregulated and 1,375 downregulated genes in rd10 relative to WT retinas (**Fig 2C**). Comparison with the previously published analysis of nuclear genes identified 1,947 common DEGs, all with concordant direction of change, including 886 upregulated and 1,061 downregulated genes. Log2 fold changes were strongly correlated between analyses (Pearson r = 0.988; Spearman ρ = 0.972), demonstrating close agreement with the previous DESeq2 analysis. Genomic annotation showed that significant DEGs were distributed across most autosomes and chromosome X (**Fig 2D**).

Expression heatmaps showed consistent group-specific patterns (**Fig 3A**), while Reactome enrichment identified immune-, interleukin-, and complement-associated pathways enriched among predominantly upregulated DEGs and phototransduction-associated pathways enriched among predominantly downregulated DEGs (**Fig 3B–C**). Detailed enriched pathways, representative differentially expressed genes, and gene– pathway relationships are provided in the Supplementary Results.

**Fig 3.**
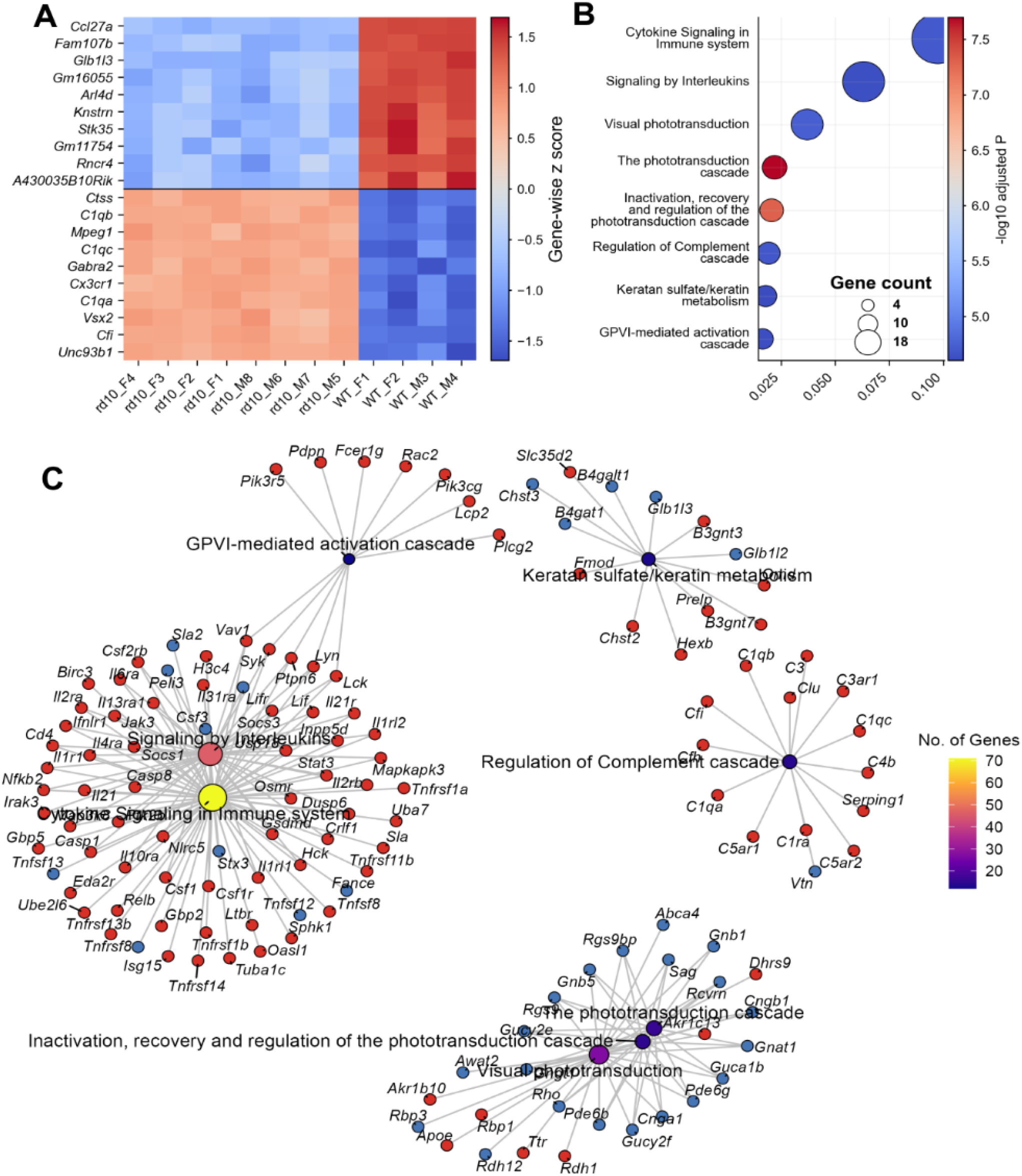
Differential-expression patterns and functional enrichment in rd10 versus WT mouse retinas using CoTRA. **(A)** Heatmap of the ten most significantly upregulated and downregulated genes; values are gene-wise Z-scores from normalized expression. **(B)** Reactome over-representation analysis of DEGs, where gene ratio is shown on the x-axis, dot size indicates DEG count, and color represents −log10 adjusted P value. **(C)** Gene–concept network linking enriched pathways to contributing DEGs; pathway-node size/color reflect associated gene number, gene-node color indicates upregulation (red) or downregulation (blue), and edges denote gene–pathway membership and shared genes.

Overall, the bulk case study demonstrates that CoTRA reproduces established rd10-associated transcriptional changes while integrating quality control, differential-expression analysis, genomic annotation, functional enrichment, and visualization within a single workflow.

### Single-cell RNA-seq case study

To demonstrate the scRNA-seq workflow, CoTRA was applied to the publicly available retinal dataset GSE234797 [41]. The analyzed dataset contained 4,478 WT and 5,952 rd10 cells. The source 10x Genomics HDF5 matrices are distributed with the CoTRA repository as test inputs, while the retained cell set and analytical parameters used for the case study are documented in Supplementary Information.

Cell-level quality-control metrics were evaluated before downstream analysis (**S3 Fig**), and 2,000 highly variable genes were selected for dimensionality reduction (**Fig 4A-B**). UMAP and t-SNE representations showed the transcriptional structure of the combined WT and rd10 dataset (**Fig 4C-D**). Graph-based clustering resolved 16 clusters (**Fig 4E-F**), cluster composition by condition is shown in (**Fig 4G**) and cell-type annotation identified the major retinal populations, including rods, cones, bipolar cells, amacrine cells, ganglion cells, Müller glia, microglia, and endothelial cells (**Fig 4H**). Cluster-associated markers and representative gene-expression patterns supporting these assignments are provided in the Supplementary Results and **S4-S5 Figs**.

**Fig 4.**
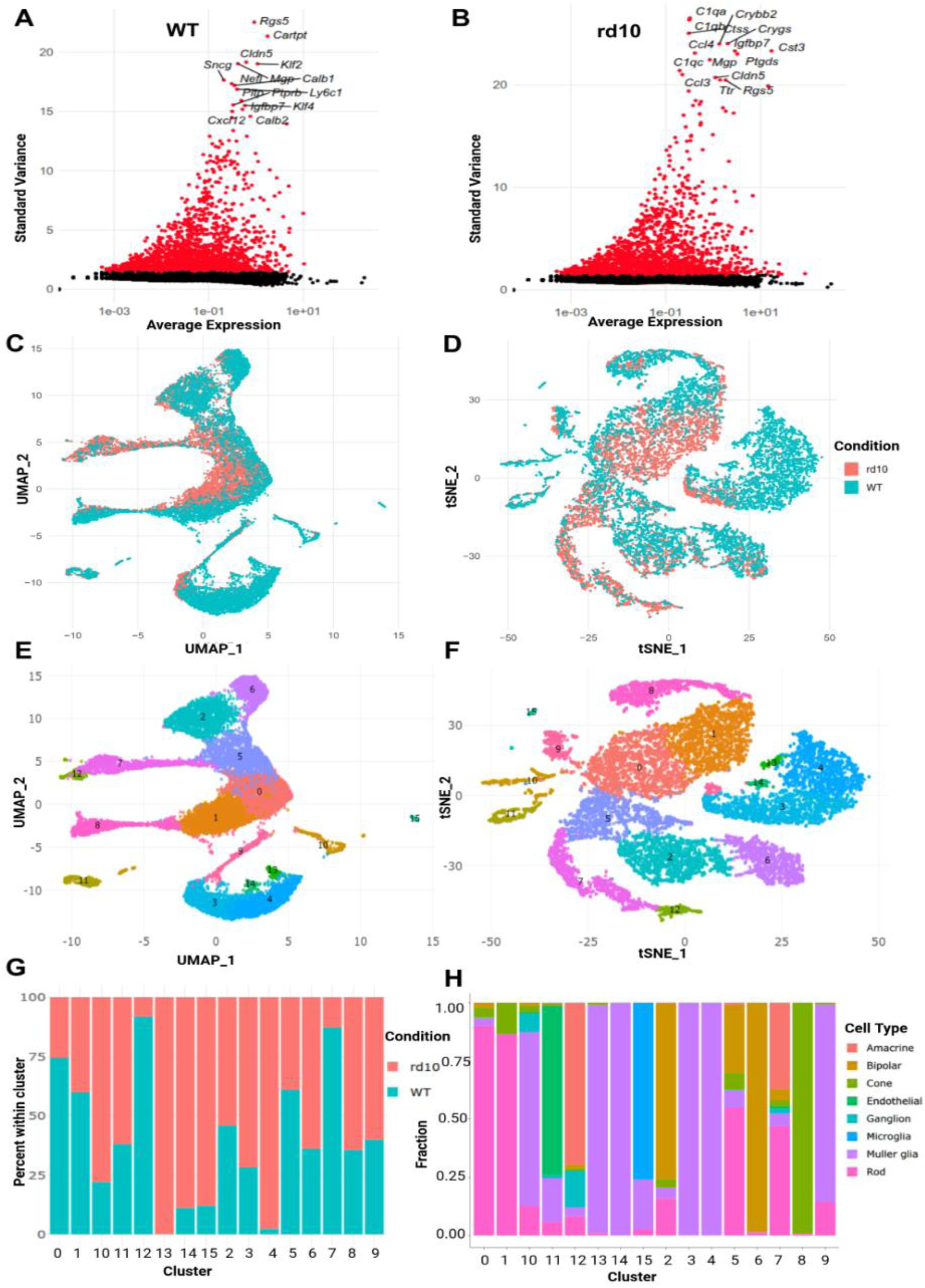
Single-cell RNA-seq analysis of WT and rd10 mouse retinal cells with CoTRA. **(A–B)** Highly variable gene analysis for WT and rd10 cells; red points indicate the 2,000 selected variable genes, black points non-variable genes, and selected high-variance genes are labeled. **(C–D)** UMAP and t-SNE embeddings colored by condition. **(E–F)** UMAP and t-SNE representations of 16 CoTRA-derived clusters (0–15). **(G)** WT/rd10 contribution within each cluster. **(H)** Annotated cell-type composition per cluster. Panels G–H demonstrate composition/differential-abundance visualization; because each condition comprised one pooled library, differences are descriptive and not statistically validated abundance changes.

Cell-type-specific differential-expression analysis was subsequently demonstrated in rod photoreceptors and Müller glia, revealing distinct rd10-versus-WT transcriptional profiles within the two populations (**Fig 5A-B**). Hallmark pathway activity was assessed using UCell to demonstrate single-cell gene-signature analysis. Rods showed comparatively modest differences, with reactive oxygen species and oxidative phosphorylation among the largest positive rd10-versus-WT changes, whereas Müller glia showed stronger differences in inflammatory and stress-associated signatures, particularly TNFα signaling via NF-κB and interferon responses (**Fig 5C-D**). Detailed differentially expressed genes, pathway rankings, and UCell-score differences are provided in the Supplementary Results.

**Fig 5.**
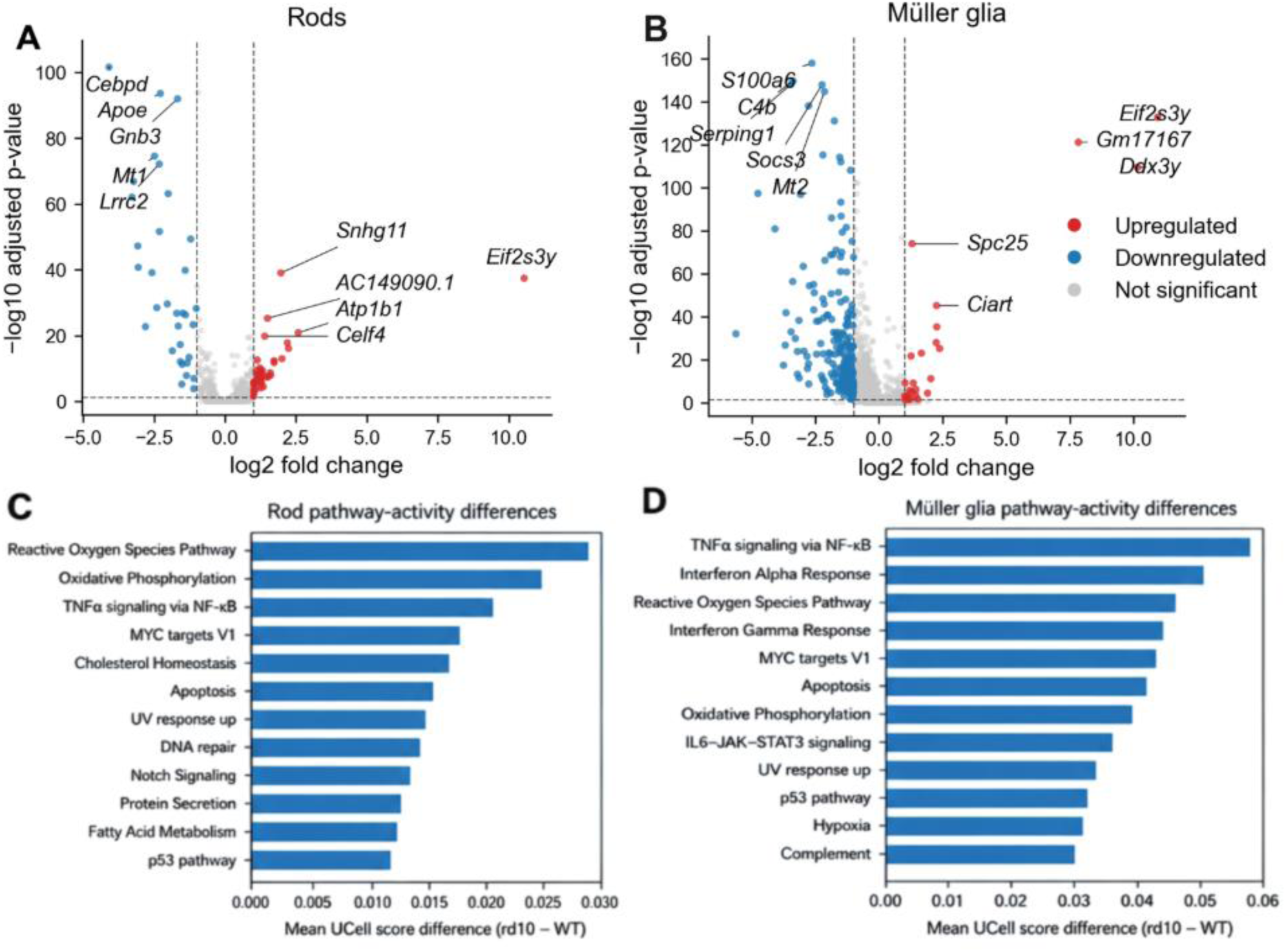
Cell-type-specific differential expression and pathway analysis in rd10 versus WT mouse retina. **(A-B)** Volcano plots for rod photoreceptors and Müller glia; each point is a gene, red/blue indicate significant differential expression in opposite directions, grey indicates genes below significance/effect-size thresholds, and selected genes are labeled. Positive and negative log2 fold changes denote opposite expression directions. **(C-D)** Ten Hallmark pathways with the largest rd10-versus-WT differences in mean UCell score for rods and Müller glia, respectively.

Because each condition was represented by a single pooled scRNA-seq library rather than independent library-level biological replicates, the cell-type-specific differential-expression, pathway-score, and composition comparisons are presented as demonstrations of CoTRA functionality and should be interpreted as descriptive condition-associated differences rather than replicate-level statistical evidence of disease effects.

### Computational performance benchmarking

For synthetic bulk RNA-seq data containing 20,000 genes, increasing the sample number from 6 to 96 increased median analytical runtime from 4.25 to 20.37 s for DESeq2 and from 1.65 to 8.74 s for edgeR, while median peak memory remained approximately 1 GB or lower (**Fig 6A-B; S3 Table**). On the real 55,291-gene, 12-sample retinal dataset, median runtimes were 12.93 s for DESeq2 and 2.53 s for edgeR, with peak memory requirements of 0.88 and 0.65 GB, respectively (**Table 1; S4 Table**). For synthetic scRNA-seq data, increasing dataset size from 2,500 to 50,000 cells increased the median combined core-workflow and marker-analysis workload from 13.29 to 299.35 s and peak memory from 0.67 to 5.10 GB (**Fig 6C-D; S5 Table**). Analysis of the complete real retinal scRNA-seq dataset containing 18,533 genes and 10,430 cells required a median of 46.68 s and 3.36 GB peak memory (**Table 1; S6 Table**). All measured runs completed successfully; step-level computational requirements are provided in **S7 Table**.

**Fig 6.**
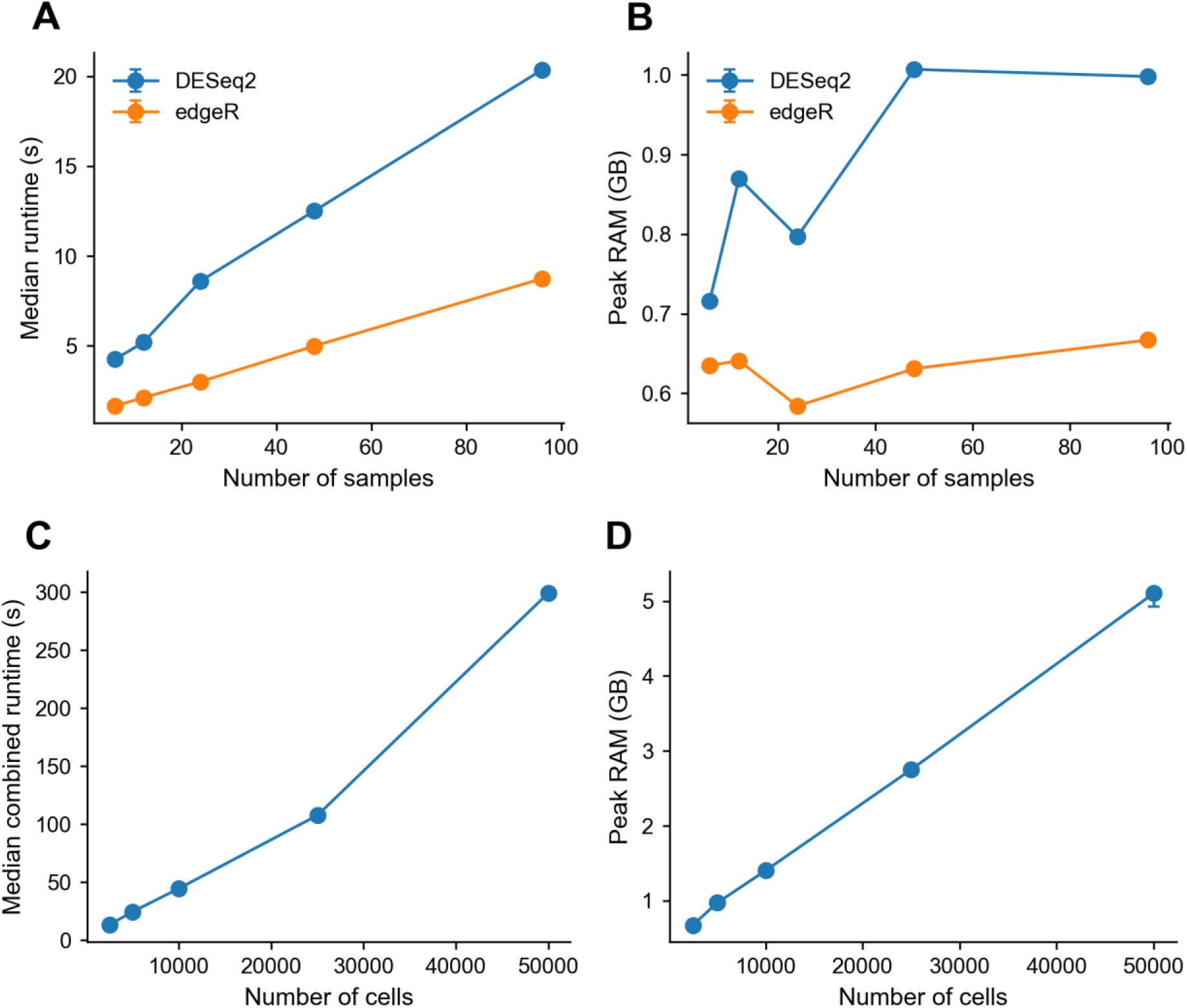
Computational performance of CoTRA across bulk and single-cell RNA-seq dataset sizes. **(A-B)** Median runtime and peak resident memory for DESeq2 and edgeR on synthetic bulk matrices with 20,000 genes and 6–96 samples. **(C-D)** Median combined core-workflow plus truth-label marker-analysis runtime and peak memory for synthetic scRNA-seq datasets with 20,000 genes and 2,500–50,000 cells. The core workflow included Seurat object/QC construction, normalization, HVG selection, scaling, PCA, UMAP, neighbor graph construction, and Louvain clustering. Marker timing used known simulated labels. Points represent the median of five single-threaded benchmark runs, and vertical bars show the interquartile range (25th–75th percentiles) across replicates. Details are in **S3, S5, and S7 Tables**.

**Table 1:** Summary of CoTRA computational performance across synthetic and real bulk and single-cell RNA-seq benchmarks.

| Dataset | Method/<br>workflow | Maximum<br>benchmark<br>size | Median<br>runtime (s) | Peak<br>RAM<br>(GB) | Hardware |
| --- | --- | --- | --- | --- | --- |
| Synthetic bulk | DESeq2 | 20,000 genes<br>× 96 samples | 20.367 | 0.998 | Ubuntu 24.04.4; Intel i9-10900; 31 GB RAM; single-thread |
| Synthetic bulk | edgeR | 20,000 genes<br>× 96 samples | 8.743 | 0.667 | Ubuntu 24.04.4; Intel i9-10900; 31 GB RAM; single-thread |
| Real retinal bulk | DESeq2 | 55,291 genes<br>× 12 samples | 12.929 | 0.881 | Ubuntu 24.04.4; Intel i9-10900; 31 GB RAM; single-thread |
| Real retinal bulk | edgeR | 55,291 genes<br>× 12 samples | 2.525 | 0.652 | Ubuntu 24.04.4; Intel i9-10900; 31 GB RAM; single-thread |
| Synthetic scRNA-seq | Core + truth-label marker workload | 20,000 genes<br>× 50,000 cells | 299.354 | 5.104 | Ubuntu 24.04.4; Intel i9-10900; 31 GB RAM; single-thread |
| Real retinal scRNA-seq | Core + inferred-cluster markers | 18,533 genes<br>× 10,430 cells | 46.675 | 3.362 | Ubuntu 24.04.4; Intel i9-10900; 31 GB RAM; single-thread |

### Reproducibility and implementation validation

CoTRA reproduced the corresponding direct scripted implementations when identical inputs and analytical settings were used. For bulk RNA-seq, DESeq2 and edgeR produced identical tested and significant gene sets, significant-set Jaccard indices of 1.000, 100% directional concordance, and log2 fold-change correlations of 1.000 (**S8 Table**). For scRNA-seq, CoTRA and direct Seurat selected the same 2,000 highly variable genes, produced perfectly concordant PCA representations for PCs 1-7, and assigned all 15,106 cells to the same 16 clusters (ARI = 1.000; NMI = 1.000). All 38,491 marker gene cluster pairs were also reproduced, with marker-set Jaccard and log2 fold-change correlations of 1.000 (**S9 Table**). These results demonstrate that the CoTRA graphical workflows reproduce their corresponding scripted implementations under matched input data and parameter settings.

### Comparison with existing transcriptomic analysis platforms

CoTRA was compared with 14 representative transcriptomic analysis platforms across 49 predefined criteria. CoTRA provided broad coverage across both bulk and single-cell RNA-seq workflows within a single local R/Shiny environment, including configurable differential-expression analysis, functional interpretation, clustering, annotation, trajectory analysis, pathway activity, cell-cell communication, and export-oriented reporting. The comparison also identified areas in which individual alternative platforms provided capabilities not currently implemented in CoTRA. The complete criterion-level comparison is provided in **S11 Table**, and the overall feature distribution is summarized in **S2 Fig**.

## AVAILABILITY AND FUTURE DIRECTIONS

CoTRA version 1.0.0 is available as an open-source R package from the public GitHub repository https://github.com/UmairSeemab/CoTRA under the GPL-3 license. The version corresponding to this publication is permanently archived on Zenodo (doi:10.5281/zenodo.22643667). The package requires R ≥ 4.4.0 and can be executed locally through its Shiny interface on Linux, Windows, and macOS. The repository provides source code, documentation, representative bulk and single-cell RNA-seq test datasets, and reproducibility resources underlying the computational-performance and implementation-validation analyses. Detailed installation instructions, test-data specifications, expected outputs, and reproduction procedures are provided in the Supplementary Information and repository. Software issues, feature requests, and contributions can be submitted through the GitHub issue tracker.

CoTRA currently focuses on downstream transcriptomic analysis and does not perform FASTQ preprocessing, read alignment, or joint integration of bulk and single-cell RNA-seq datasets. Computational requirements for large scRNA-seq datasets remain influenced by the underlying R and Seurat implementations, and inferential analyses remain dependent on appropriate experimental design and biological replication. Future development will focus on improving single-cell scalability, strengthening multi-sample and batch-aware analyses, expanding validated annotation and downstream-analysis options, and enhancing automated workflow provenance and reproducibility reporting.

## Acknowledgments

The authors thank Dr. Nicholas Bariesheff for carefully proof-reading the manuscript and providing valuable feedback.

## Supporting Information

**S1 Fig. CoTRA graphical user interface.** Representative CoTRA interface showing workflow navigation, user-configurable analytical parameters, visualization of results, download options, and integrated interpretation guidance.

**S2 Fig. Feature comparison of CoTRA with representative graphical RNA-seq analysis platforms.** Heatmap comparing CoTRA and 14 other platforms across 49 predefined criteria covering deployment, usability, input handling, bulk and single-cell RNA-seq workflows, visualization, biological interpretation, advanced analyses, export, reporting, and reproducibility.

**S3 Fig. Quality-control characteristics of WT and rd10 mouse retinal single-cell RNA-seq data.** Violin plots showing distributions of detected genes per cell, total RNA counts, mitochondrial transcript percentage, and ribosomal transcript percentage.

**S4 Fig. Cluster-associated marker gene expression in the CoTRA analysis.** Dot plot showing the three highest-ranking marker genes for each of the 16 identified clusters; dot size represents the proportion of expressing cells and color represents average expression.

**S5 Fig. UMAP visualization of representative cluster-associated genes.** Feature plots showing expression distributions of representative retinal marker genes across the UMAP embedding.

**S1 Table. CoTRA analysis methods, default implementations, study settings, and user-configurable parameters.**

**S2 Table. Computational environment used for CoTRA benchmarking.**

**S3 Table. Synthetic bulk RNA-seq computational benchmark results.**

**S4 Table. Real retinal bulk RNA-seq computational benchmark results.**

**S5 Table. Synthetic scRNA-seq computational benchmark results.**

**S6 Table. Real retinal scRNA-seq computational benchmark results.**

**S7 Table. Step-level computational timing results.**

**S8 Table. Concordance between CoTRA and direct scripted bulk differential-expression implementations.** Concordance results for DESeq2 and edgeR analyses performed using identical input data and analytical settings.

**S9 Table. Concordance between CoTRA and direct scripted Seurat analysis.** Concordance of highly variable genes, PCA coordinates, cluster assignments, and marker-analysis results under matched input data and parameters.

**S10 Table. CoTRA test datasets, fixed validation settings, and expected outputs.** Characteristics of the supplied bulk and scRNA-seq test datasets together with the fixed settings and expected outputs used for reproducibility checks.

**S11 Table. Feature comparison of CoTRA and 14 representative transcriptomic analysis platforms across 49 predefined criteria.** Includes the criterion-level comparison, definitions, evidence sources, software versions or releases, and evaluation information.

**S1 Data. Run-level measured computational-benchmark outputs.** Run-level runtime, step-level timing, and GNU time -v resource measurements underlying the reported computational performance results.

**S2 Data. Processed computational benchmark summaries and figure-source data.** Machine-readable benchmark summaries, computational-environment information, step-level summaries, and source data used to generate the computational-performance figure.

**S3 Data. Preliminary computational benchmark viability runs.** Preliminary runs performed to confirm successful execution before the five measured replicates; these runs were excluded from all reported benchmark statistics.

**S4 Data. Machine-readable implementation-validation outputs.** Bulk and scRNA-seq concordance summaries, run-level comparisons, PCA concordance results, cluster contingency matrices, and cluster-label mappings underlying S8 and S9 Tables.

**S1 Code. CoTRA computational benchmarking scripts.** Scripts used for synthetic-data generation, real-data preparation, computational benchmark execution, result summarization, and computational-performance figure generation.

**S2 Code. CoTRA implementation-validation scripts.** Scripts used to compare CoTRA with direct DESeq2, edgeR, and Seurat implementations using matched input data and analytical settings.

## REFERENCES

1. Mortazavi A, Williams BA, McCue K, Schaeffer L, Wold B. Mapping and quantifying mammalian transcriptomes by RNA-Seq. Nat Methods. 2008;5: 621– 628. doi:10.1038/nmeth.1226

2. Tang F, Barbacioru C, Wang Y, Nordman E, Lee C, Xu N, et al. mRNA-Seq whole-transcriptome analysis of a single cell. Nat Methods. 2009;6: 377–382. doi:10.1038/nmeth.1315

3. Conesa A, Madrigal P, Tarazona S, Gomez-Cabrero D, Cervera A, McPherson A, et al. A survey of best practices for RNA-seq data analysis. Genome Biol. 2016;17: 13. doi:10.1186/s13059-016-0881-8

4. Lähnemann D, Köster J, Szczurek E, McCarthy DJ, Hicks SC, Robinson MD, et al. Eleven grand challenges in single-cell data science. Genome Biol. 2020;21: 31. doi:10.1186/s13059-020-1926-6

5. Lun ATL, Chen Y, Smyth GK. It’s DE-licious: A Recipe for Differential Expression Analyses of RNA-seq Experiments Using Quasi-Likelihood Methods in edgeR. 2016. pp. 391–416. doi:10.1007/978-1-4939-3578-9_19

6. Love MI, Huber W, Anders S. Moderated estimation of fold change and dispersion for RNA-seq data with DESeq2. Genome Biol. 2014;15: 550. doi:10.1186/s13059-014-0550-8

7. Robinson MD, McCarthy DJ, Smyth GK. *edgeR*: a Bioconductor package for differential expression analysis of digital gene expression data. Bioinformatics. 2010;26: 139–140. doi:10.1093/bioinformatics/btp616

8. Hänzelmann S, Castelo R, Guinney J. GSVA: gene set variation analysis for microarray and RNA-Seq data. BMC Bioinformatics. 2013;14: 7. doi:10.1186/1471-2105-14-7

9. Wu T, Hu E, Xu S, Chen M, Guo P, Dai Z, et al. clusterProfiler 4.0: A universal enrichment tool for interpreting omics data. The Innovation. 2021;2: 100141. doi:10.1016/j.xinn.2021.100141

10. Satija R, Farrell JA, Gennert D, Schier AF, Regev A. Spatial reconstruction of single-cell gene expression data. Nat Biotechnol. 2015;33: 495–502. doi:10.1038/nbt.3192

11. Wolf FA, Angerer P, Theis FJ. SCANPY: large-scale single-cell gene expression data analysis. Genome Biol. 2018;19: 15. doi:10.1186/s13059-017-1382-0

12. Amezquita RA, Lun ATL, Becht E, Carey VJ, Carpp LN, Geistlinger L, et al. Orchestrating single-cell analysis with Bioconductor. Nat Methods. 2020;17: 137–145. doi:10.1038/s41592-019-0654-x

13. Aran D, Looney AP, Liu L, Wu E, Fong V, Hsu A, et al. Reference-based analysis of lung single-cell sequencing reveals a transitional profibrotic macrophage. Nat Immunol. 2019;20: 163–172. doi:10.1038/s41590-018-0276-y

14. Street K, Risso D, Fletcher RB, Das D, Ngai J, Yosef N, et al. Slingshot: cell lineage and pseudotime inference for single-cell transcriptomics. BMC Genomics. 2018;19: 477. doi:10.1186/s12864-018-4772-0

15. Andreatta M, Carmona SJ. UCell: Robust and scalable single-cell gene signature scoring. Comput Struct Biotechnol J. 2021;19: 3796–3798. doi:10.1016/j.csbj.2021.06.043

16. Finak G, McDavid A, Yajima M, Deng J, Gersuk V, Shalek AK, et al. MAST: a flexible statistical framework for assessing transcriptional changes and characterizing heterogeneity in single-cell RNA sequencing data. Genome Biol. 2015;16: 278. doi:10.1186/s13059-015-0844-5

17. Haque A, Engel J, Teichmann SA, Lönnberg T. A practical guide to single-cell RNA-sequencing for biomedical research and clinical applications. Genome Med. 2017;9: 75. doi:10.1186/s13073-017-0467-4

18. Attwood TK, Blackford S, Brazas MD, Davies A, Schneider MV. A global perspective on evolving bioinformatics and data science training needs. Brief Bioinform. 2019;20: 398–404. doi:10.1093/bib/bbx100

19. Rung J, Brazma A. Reuse of public genome-wide gene expression data. Nat Rev Genet. 2013;14: 89–99. doi:10.1038/nrg3394

20. Clough E, Barrett T, Wilhite SE, Ledoux P, Evangelista C, Kim IF, et al. NCBI GEO: archive for gene expression and epigenomics data sets: 23-year update. Nucleic Acids Res. 2024;52: D138–D144. doi:10.1093/nar/gkad965

21. Hulsen T, Jamuar SS, Moody AR, Karnes JH, Varga O, Hedensted S, et al. From Big Data to Precision Medicine. Front Med (Lausanne). 2019;6. doi:10.3389/fmed.2019.00034

22. Athar A, Füllgrabe A, George N, Iqbal H, Huerta L, Ali A, et al. ArrayExpress update – from bulk to single-cell expression data. Nucleic Acids Res. 2019;47: D711–D715. doi:10.1093/nar/gky964

23. Byrd JB, Greene AC, Prasad DV, Jiang X, Greene CS. Responsible, practical genomic data sharing that accelerates research. Nat Rev Genet. 2020;21: 615– 629. doi:10.1038/s41576-020-0257-5

24. Fillbrunn A, Dietz C, Pfeuffer J, Rahn R, Landrum GA, Berthold MR. KNIME for reproducible cross-domain analysis of life science data. J Biotechnol. 2017;261: 149–156. doi:10.1016/j.jbiotec.2017.07.028

25. Sandve GK, Nekrutenko A, Taylor J, Hovig E. Ten Simple Rules for Reproducible Computational Research. PLoS Comput Biol. 2013;9: e1003285. doi:10.1371/journal.pcbi.1003285

26. Cohen-Boulakia S, Belhajjame K, Collin O, Chopard J, Froidevaux C, Gaignard A, et al. Scientific workflows for computational reproducibility in the life sciences: Status, challenges and opportunities. Future Generation Computer Systems. 2017;75: 284–298. doi:10.1016/j.future.2017.01.012

27. Afgan E, Galaxy Community. The Galaxy platform for accessible, reproducible and collaborative biomedical analyses: 2022 update. Nucleic Acids Res. 2022;50: W345–W351. doi:10.1093/nar/gkac247

28. Ge SX, Son EW, Yao R. iDEP: an integrated web application for differential expression and pathway analysis of RNA-Seq data. BMC Bioinformatics. 2018;19:534. doi:10.1186/s12859-018-2486-6

29. Torre D, Lachmann A, Ma’ayan A. BioJupies: Automated Generation of Interactive Notebooks for RNA-Seq Data Analysis in the Cloud. Cell Syst. 2018;7: 556–561.e3. doi:10.1016/j.cels.2018.10.007

30. Kucukural A, Yukselen O, Ozata DM, Moore MJ, Garber M. DEBrowser: interactive differential expression analysis and visualization tool for count data. BMC Genomics. 2019;20: 6. doi:10.1186/s12864-018-5362-x

31. Marini F, Linke J, Binder H. ideal: an R/Bioconductor package for interactive differential expression analysis. BMC Bioinformatics. 2020;21: 565. doi:10.1186/s12859-020-03819-5

32. Su S, Law CW, Ah-Cann C, Asselin-Labat M-L, Blewitt ME, Ritchie ME. Glimma: interactive graphics for gene expression analysis. Bioinformatics. 2017;33: 2050– 2052. doi:10.1093/bioinformatics/btx094

33. Sun L, Dong S, Ge Y, Fonseca JP, Robinson ZT, Mysore KS, et al. DiVenn: An Interactive and Integrated Web-Based Visualization Tool for Comparing Gene Lists. Front Genet. 2019;10. doi:10.3389/fgene.2019.00421

34. Stuart T, Butler A, Hoffman P, Hafemeister C, Papalexi E, Mauck WM, et al. Comprehensive Integration of Single-Cell Data. Cell. 2019;177: 1888–1902.e21. doi:10.1016/j.cell.2019.05.031

35. Rue-Albrecht K, Marini F, Soneson C, Lun ATL. iSEE: Interactive SummarizedExperiment Explorer. F1000Res. 2018;7: 741. doi:10.12688/f1000research.14966.1

36. Aussel R, Asif M, Chenag S, Jaeger S, Milpied P, Spinelli L. ShIVA: a user-friendly and interactive interface giving biologists control over their single-cell RNA-seq data. Sci Rep. 2023;13: 14377. doi:10.1038/s41598-023-40959-z

37. Puente-Santamaría L, del Peso L. SinglePointRNA, an user-friendly application implementing single cell RNA-seq analysis software. PLoS One. 2024;19: e0300567. doi:10.1371/journal.pone.0300567

38. Dimitrov D, Gu Q. BingleSeq: a user-friendly R package for bulk and single-cell RNA-Seq data analysis. PeerJ. 2020;8: e10469. doi:10.7717/peerj.10469

39. Weber C, Hirst MB, Ernest B, Schaub NJ, Wilson KM, Wang K, et al. SEQUIN is an R/Shiny framework for rapid and reproducible analysis of RNA-seq data. Cell Reports Methods. 2023;3: 100420. doi:10.1016/j.crmeth.2023.100420

40. Monier B, McDermaid A, Wang C, Zhao J, Miller A, Fennell A, et al. IRIS-EDA: An integrated RNA-Seq interpretation system for gene expression data analysis. PLoS Comput Biol. 2019;15: e1006792. doi:10.1371/journal.pcbi.1006792

41. Leinonen H, Zhang J, Occelli LM, Seemab U, Choi EH, L.P. Marinho LF, et al. A combination treatment based on drug repurposing demonstrates mutation-agnostic efficacy in pre-clinical retinopathy models. Nat Commun. 2024;15: 5943. doi:10.1038/s41467-024-50033-5

