## Supplementary Text for "CoTRA: a comprehensive R/Shiny framework for transparent bulk and single-cell RNA-seq analysis"

S1 Fig. CoTRA Graphical User Interface (GUI).

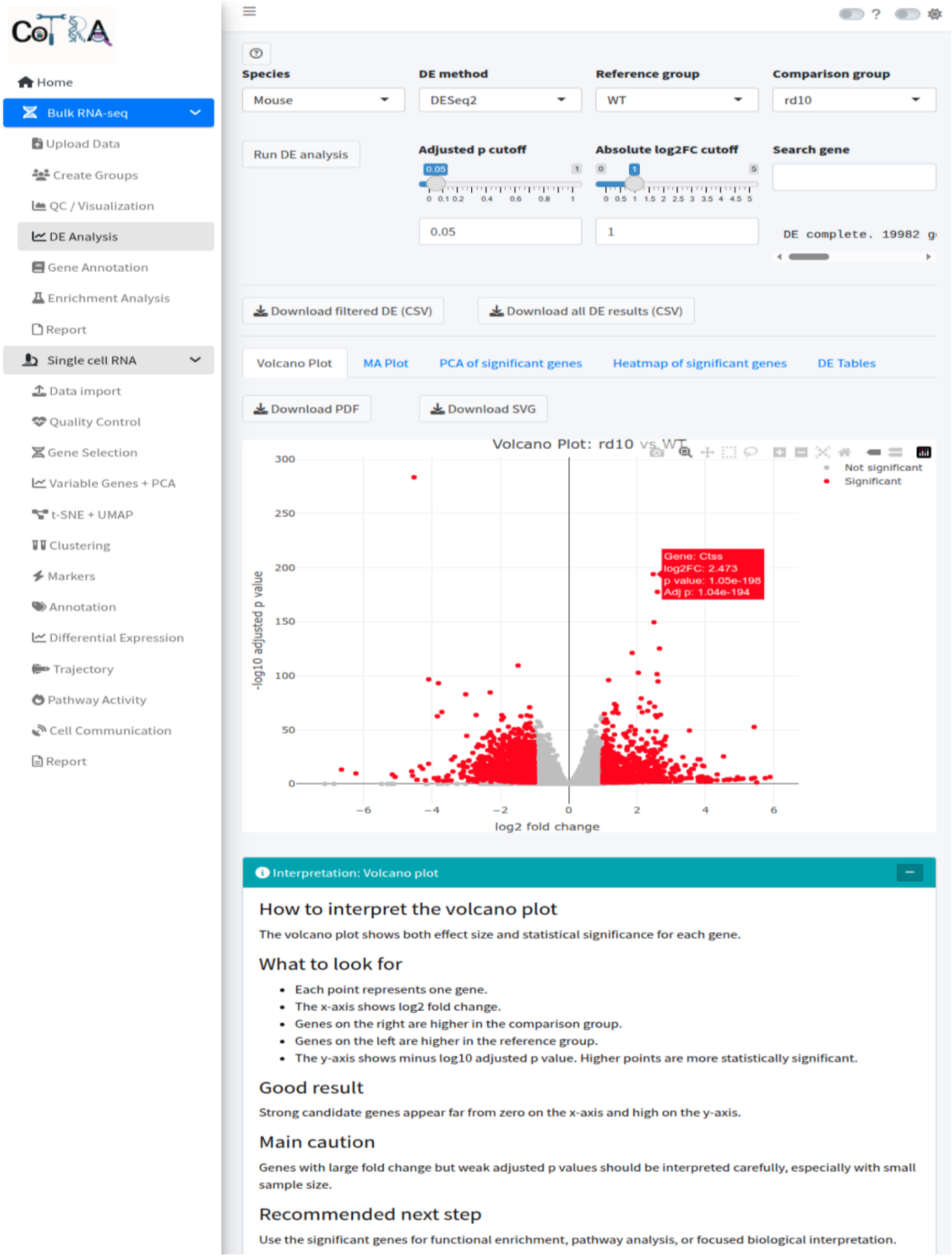

**S1 Fig. CoTRA graphical user interface.** Representative CoTRA graphical user interface showing workflow navigation, user-configurable analytical parameters, visualization of results and download options, as well as integrated interpretation guidance. The displayed bulk RNA-seq differential expression module illustrates the selection of species, differential expression method, reference and comparison groups, adjusted P-value and log2 fold-change thresholds, gene search, visualization tabs, and downloadable outputs.

S2 Fig. Feature comparison of CoTRA.

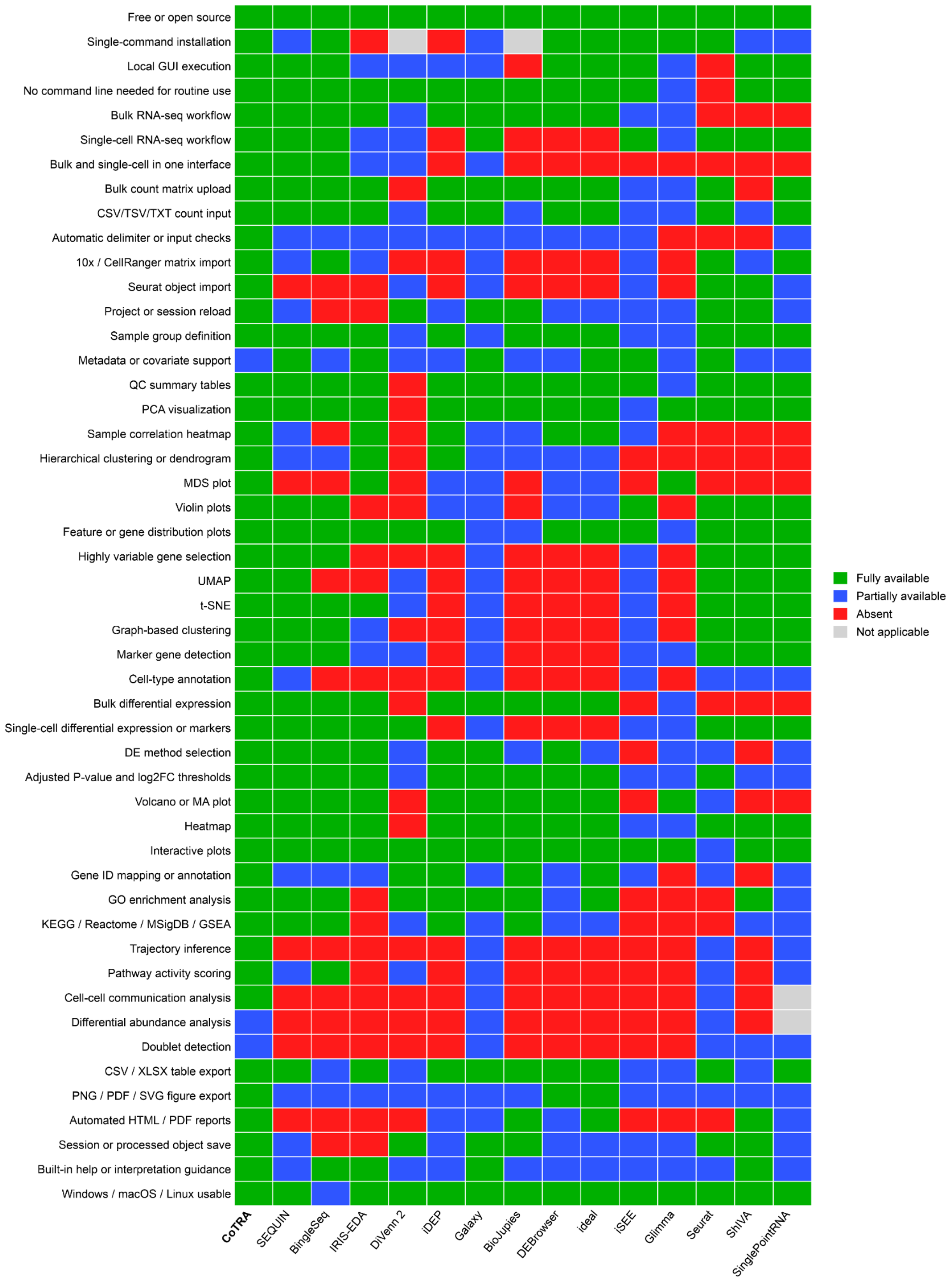

**S2 Fig. Feature comparison of CoTRA with representative graphical RNA-seq analysis platforms.** Heatmap comparing CoTRA and 14 existing tools across 49 predefined criteria covering installation and usability, data input, quality control, bulk and single-cell RNA-seq workflows, differential expression, visualization, functional interpretation, advanced single-cell analyses, export, reporting, and reproducibility. Features were classified as **fully available** (green), **partially available** (blue), **absent** (red), or **not**

**applicable** (grey). Detailed criteria, classifications, evidence sources, software versions, and evaluation information are provided in the S11 Table.

**SUPPLEMENTARY RESULTS**

### Supplementary bulk RNA-seq case-study results

Differential-expression analysis was performed using DESeq2 for the rd10-versus-WT comparison. The filtering, differential-expression, significance, and downstream-analysis settings used for the bulk case study are summarized in S1 Table. Genes were retained for statistical testing according to the count-filtering procedure implemented in CoTRA, and differential-expression significance was defined using an adjusted P value (Benjamini–Hochberg) < 0.05 together with an absolute log2 fold change ≥ 1. Genes meeting both criteria with log2 fold change ≥ 1 were classified as upregulated in rd10 relative to WT, whereas genes with log2 fold change ≤ -1 were classified as downregulated. Genes not meeting both the adjusted P-value and effect-size criteria were not included in the significant DEG set used for downstream interpretation. Using these criteria, CoTRA identified 2,518 significant DEGs, comprising 1,143 upregulated and 1,375 downregulated genes.

Detailed inspection of the differential-expression results identified prominent transcriptional changes between rd10 and WT mouse retinas. Among the highly significant upregulated genes were *Ctss, C1qb, Mpeg1, C1qc,* and *Gabra2*, whereas *Ccl27a, Fam107b, Glb1l3, Gm16055,* and *Arl4d* were among the strongly downregulated genes. Increased expression of immune- and microglia-associated genes, including *Ctss, C1qa, C1qb, C1qc, Mpeg1,* and *Cx3cr1*, was evident in rd10 retinas. A heatmap containing the ten most significantly upregulated and ten most significantly downregulated genes further showed consistent group-specific expression patterns across individual samples, supporting the separation observed in the sample-level correlation and principal-component analyses.

Reactome over-representation analysis was used to characterize biological processes associated with significant differentially expressed genes. The enrichment input therefore consisted only of genes meeting the adjusted P value < 0.05 and |log2 fold change| ≥ 1 criteria defined above. Pathway significance was assessed using multiple-testing-adjusted P values. Enriched pathways included Cytokine Signaling in Immune System, Signaling by Interleukins, Visual phototransduction, The phototransduction cascade, Inactivation, recovery and regulation of the phototransduction cascade, Regulation of Complement cascade, keratan sulfate/keratin metabolism, and GPVI-mediated activation cascade. Because over-representation analysis does not estimate the direction of pathway regulation, directional interpretation was based on the expression direction of the DEGs contributing to each enriched pathway. Immune-, interleukin-, and complement-associated pathways contained predominantly upregulated genes in rd10 retinas, whereas phototransduction-related pathways contained predominantly downregulated genes. These patterns were consistent with increased inflammatory and complement-associated transcription together with reduced photoreceptor-associated expression in the degenerating retina. In the enrichment dot plot, gene ratio represents the proportion of DEGs associated with each pathway, dot size represents the number of contributing DEGs, and color represents $-\log_{10}$ of the adjusted P-value.

The gene–pathway network provided additional information on the relationships between enriched pathways and their contributing genes. Cytokine Signaling in Immune System, Signaling by Interleukins, and Regulation of Complement cascade formed interconnected modules involving predominantly upregulated genes such as *C1qa, C1qb, C1qc, Clu, C3, Csf1r, Jak3,* and *Stat3*. In contrast, Visual phototransduction and related phototransduction pathways were linked predominantly to downregulated photoreceptor-associated genes including *Rho, Pde6g, Guca1b, Gngt1, Cngb1,* and *Sag*. Several genes were shared among biologically related pathways, illustrating how CoTRA's gene concept network visualization can identify common molecular components underlying multiple enrichment signals. Accordingly, descriptions of pathways associated with increased or decreased expression refer to the direction of their contributing DEGs rather than to directional statistic produced by the over-representation analysis itself.

Together, these detailed results complement the main-text analysis by showing that CoTRA can move from sample-level quality assessment and differential-expression testing to gene-level visualization, pathway enrichment, and network-based interpretation within the same bulk RNA-seq workflow.

### Supplementary single-cell RNA-seq case-study results

The retinal scRNA-seq case study used the publicly available GSE234797 dataset and focused on WT and rd10 retinal cells. The analyzed dataset contained 4,478 WT cells and 5,952 rd10 cells. Before downstream analysis, cell-level quality-control metrics were inspected, including the number of detected genes per cell, total RNA/UMI counts, mitochondrial transcript percentage, and ribosomal transcript percentage. The distributions of these quality-control metrics are shown in **S3 Fig.** The quality-control, preprocessing, clustering, annotation, differential-expression, and pathway-analysis settings used for the case study are summarized in **S1 Table**.

Highly variable feature analysis identified 2,000 variable genes for downstream dimensionality reduction (**Fig 4A-B**). Following preprocessing, UMAP and t-SNE embeddings showed the overall transcriptional structure of the combined dataset, with substantial overlap between conditions together with regions differing in the relative contribution of WT and rd10 cells (**Fi 4C-D**). Graph-based clustering resolved 16 clusters, numbered 0–15 (**Fig 4E-F**). Cluster-associated marker genes are shown in **S4 Fig**, while representative gene-expression patterns across the low-dimensional embedding are shown in **S5 Fig**. These results further illustrated the transcriptional heterogeneity of the dataset and supported the subsequent cell-type annotation.

Cell-type annotation identified the major retinal populations represented in the dataset, including rod and cone photoreceptors, bipolar cells, amacrine cells, ganglion cells, Müller glia, microglia, and endothelial cells (**Fig 4H; S1 Table**). The relative representation of WT and rd10 cells varied among clusters (**Fig 4G**). Because each condition was represented by a single pooled scRNA-seq library rather than independent biological replicates, these differences in cluster composition were treated as descriptive and were not interpreted as statistically validated differential-abundance effects.

Cell-type-specific differential-expression analysis was demonstrated in rod photoreceptors and Müller glia (**Fig 5 A-B**). Rods showed prominent differential-expression signals including *Apoe, Grb3,* and *Mt2*, whereas Müller glia showed a broader transcriptional response involving genes including *Serping1, Socs3, Mt2, Spc25,* and *Ciart*. These analyses illustrate that CoTRA can subset annotated retinal populations and perform within-cell-type expression comparisons within the same scRNA-seq workflow. As for the cluster-composition analysis, the absence of independent library-level biological replicates means that these cell-level comparisons should be interpreted as descriptive condition-associated differences rather than replicate-level statistical evidence of disease effects.

Hallmark gene-set activity was evaluated using UCell to demonstrate pathway- and signature-level analysis at single-cell resolution. Mean pathway scores were summarized separately for selected retinal cell populations. In rod photoreceptors, the largest positive rd10-versus-WT differences included reactive oxygen species (ΔUCell = 0.028), oxidative phosphorylation (0.023), TNFα signaling via NF-κB (0.019), MYC targets V1 (0.016), cholesterol homeostasis (0.015), apoptosis (0.013), and DNA repair (0.013) (**Fig 5C**). The overall magnitude of these differences was comparatively modest but directionally coherent across several stress- and metabolism-associated signatures.

Müller glia showed larger descriptive rd10-versus-WT pathway-score differences. Increased mean UCell scores were observed for TNFα signaling via NF-κB, interferon-α response, reactive oxygen species, interferon-γ response, MYC targets V1, apoptosis, oxidative phosphorylation, IL6–JAK–STAT3 signaling, hypoxia, p53 pathway, and complement signatures (**Fig 5D**). The largest mean difference was observed for TNFα signaling via NF-κB (ΔUCell = 0.056), followed by interferon-α response (0.050), reactive oxygen species (0.045), and interferon-γ response (0.044). These results demonstrate that CoTRA can summarize pathway-level transcriptional activity within selected annotated cell populations in addition to gene-level differential-expression analysis.

The cell-type-specific differential-expression, pathway-score, and composition results reported here were generated primarily to demonstrate the functionality of the corresponding CoTRA modules. Because the WT and rd10 conditions were each represented by a single pooled scRNA-seq library, these analyses do not provide replicate-level statistical inference regarding disease-associated changes in gene expression, pathway activity, or cell abundance. Instead, they provide a reproducible demonstration of how CoTRA can progress from quality control and dimensionality reduction to clustering, cell-type annotation, within-population expression analysis, and pathway-level interpretation within a single scRNA-seq workflow.

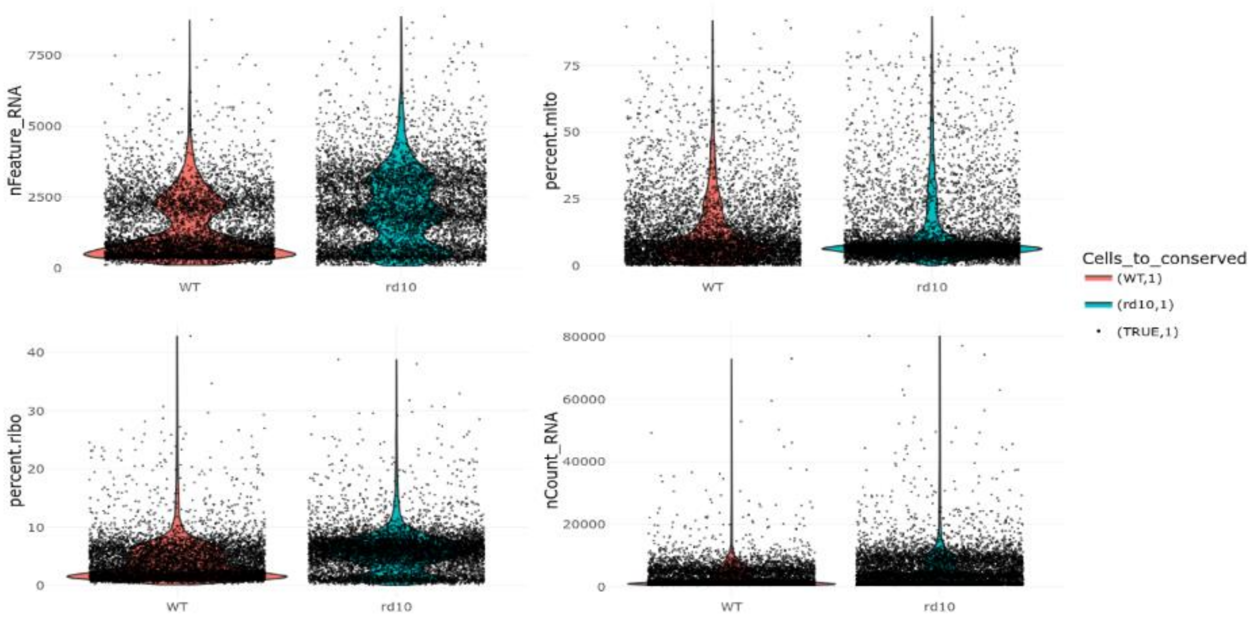

**S3 Fig. Quality control characteristics of WT and rd10 mouse retinal single-cell RNA-seq data.** Violin plots show the distributions of the number of detected genes per cell (nFeature_RNA), mitochondrial transcript percentage (percent.mito), ribosomal transcript percentage (percent.ribo), and total RNA counts per cell (nCount_RNA) in WT and rd10 mouse retinal cells. Individual points represent cells. These quality control metrics were inspected before normalization, feature selection, dimensionality reduction, and clustering.

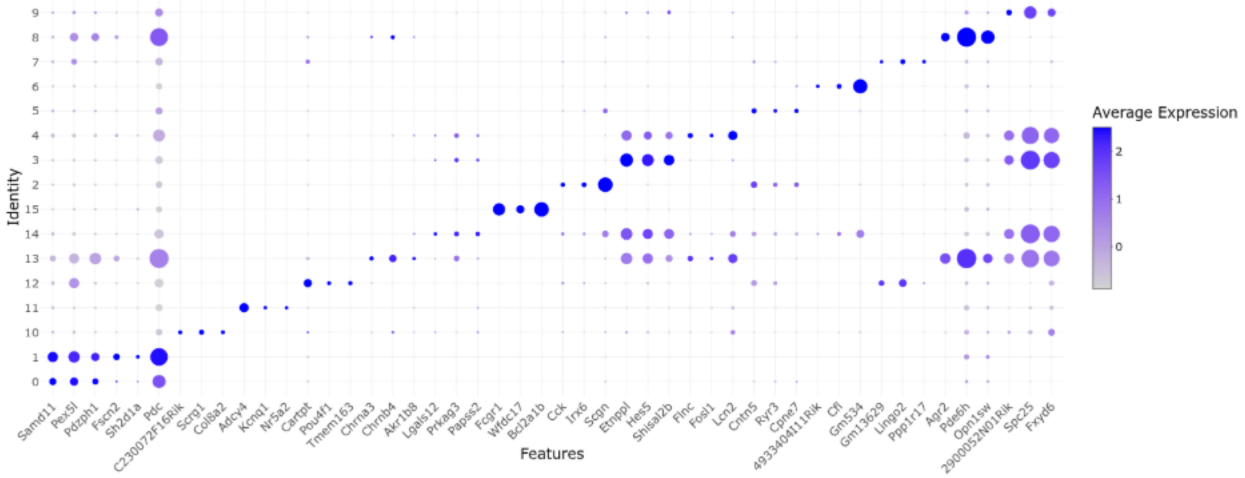

**S4 Fig. Cluster-associated marker gene expression in the CoTRA analysis.** Dot plot showing the three highest ranking marker genes identified for each of the 16 clusters. Clusters are shown on the y-axis and marker genes on the x-axis. Dot size represents the proportion of cells expressing the corresponding gene, whereas dot color represents average expression. Cluster-associated marker profiles were used to characterize transcriptional differences among clusters and support subsequent biological interpretation and cell-type annotation.

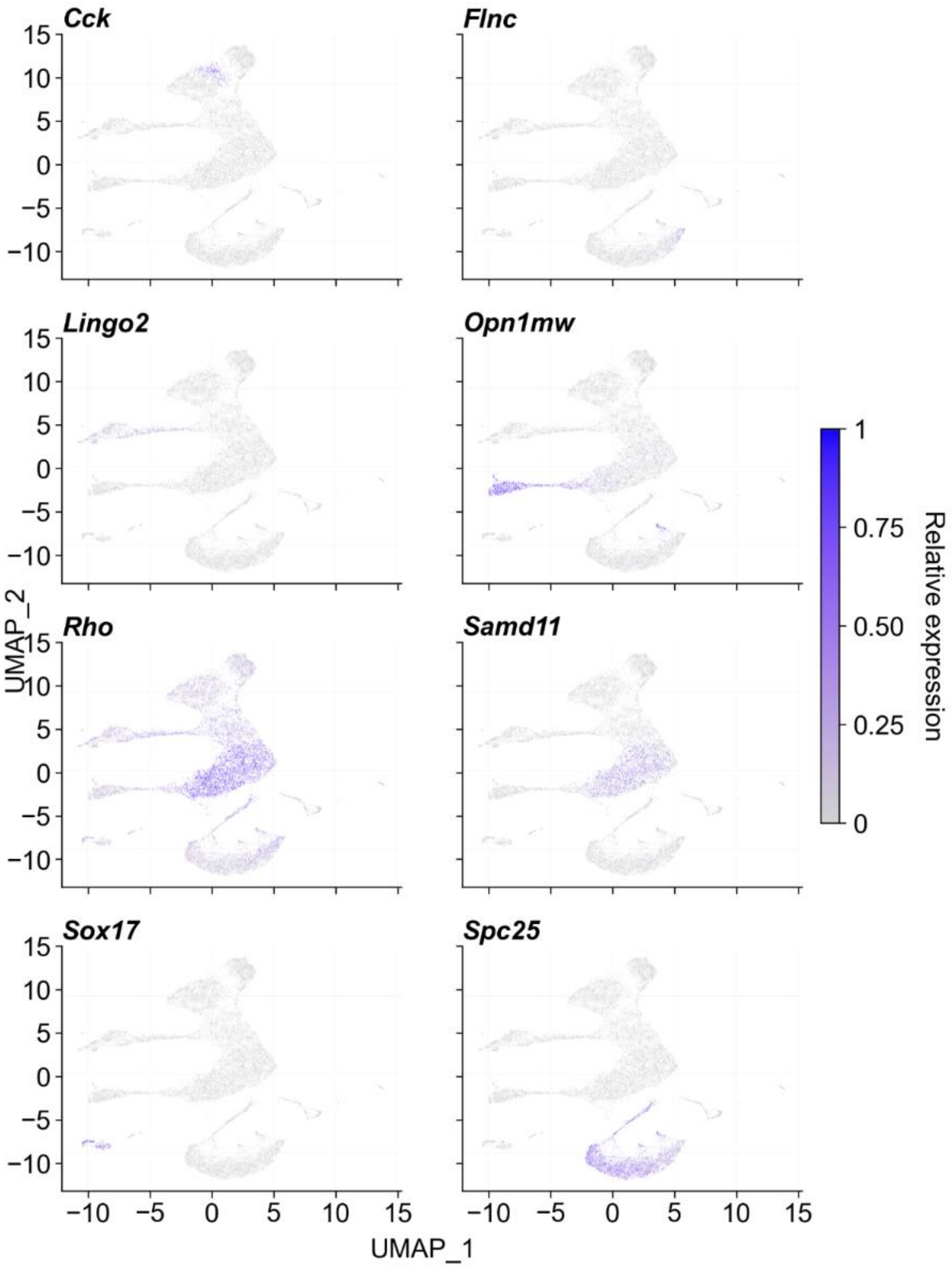

**S5 Fig. UMAP visualization of representative cluster-associated genes.** Feature plots show the expression distribution across the UMAP embedding of *Cck, Flnc, Lingo2, Opn1mw, Rho, Samd11, Sox17,* and *Spc25* across UMAP embedding. Expression values are displayed using a common relative-expression scale from 0 to 1, with increasing color intensity indicating higher relative expression. Spatially restricted expression of these representative genes provides additional visualization of cluster-associated transcriptional heterogeneity.

### SUPPLEMENTARY COMPUTATIONAL PERFORMANCE BENCHMARKING

#### Benchmark environment and measurement strategy

Computational performance was evaluated on a workstation running Ubuntu 24.04.4 LTS and equipped with an Intel Core i9-10900 processor with 10 physical cores and 20 logical processors and 31 GB RAM. The software environment included R 4.6.1, Seurat 5.5.1, SeuratObject 5.4.0, Matrix 1.7.6, DESeq2 1.52.0, and edgeR 4.10.4. OMP, OpenBLAS, MKL, VECLIB, and NUMEXPR thread counts were restricted to one where applicable to minimize variability associated with parallel execution. The complete computational environment, including BLAS/LAPACK configuration and thread restrictions, is provided in **S2 Table**.

Each dataset size and analysis configuration was evaluated in five independent measured R processes. A separate preliminary viability run was performed before the measured replicates to confirm successful execution and was excluded from all reported statistics. Synthetic datasets were generated before benchmarking and stored as fixed inputs so that the five measured replicates within each condition analyzed identical data. Random seeds were fixed where applicable.

Analytical runtime was measured around the corresponding computational workflow and excluded R startup, package loading, synthetic data generation, benchmark-input preparation, and input-file loading. Shiny user interaction, browser rendering, download operations, and automated report generation were not included. Peak resident memory represented the maximum memory footprint of the complete R process and was obtained from the maximum resident set size reported by GNU /usr/bin/time -v.

For each condition, runtime and memory measurements are summarized using the median and interquartile range (Q1–Q3). Run-to-run variability was additionally characterized using the coefficient of variation (CV),

$$\mathrm{CV}(\%) = \frac{s}{\bar{x}} \times 100$$

where s is the standard deviation and $\bar{x}$ is the arithmetic mean across the five measured replicates.

Run level CSV outputs and GNU time resource measurements underlying the reported benchmarks are provided in **S1 Data**. Processed benchmark summaries and figure-source data are provided in **S2 Data**. Benchmark generation, execution, summarization, and figure generation scripts are provided in **S1 Code**. Preliminary viability runs excluded from the reported statistics are provided separately in **S3 Data**.

#### Synthetic bulk RNA-seq benchmark

Synthetic bulk RNA-seq matrices contained 20,000 genes and balanced two-group designs with 6, 12, 24, 48, or 96 samples. Fixed benchmark matrices were generated before the measured runs and reused across all five replicates for each dataset size. DESeq2 and edgeR were evaluated independently using the corresponding implementations available through CoTRA. Generation, execution, and summarization scripts are provided in **S1 Code**.

For DESeq2, median analytical runtimes were 4.25, 5.20, 8.61, 12.52, and 20.37 s for 6, 12, 24, 48, and 96 samples, respectively. Corresponding median peak memory requirements were approximately 0.72, 0.87, 0.80, 1.01, and 1.00 GB.

For edgeR, median analytical runtimes were 1.65, 2.12, 3.00, 4.99, and 8.74 s across the same dataset sizes, while median peak-memory requirements remained approximately 0.58–0.67 GB. These values characterize computational resource requirements and should not be interpreted as a comparison of statistical accuracy between DESeq2 and edgeR.

Complete median, Q1, Q3, peak memory, process level, and successful run statistics are provided in **S3 Table**, with corresponding step-level timings in **S7 Table**.

#### Real bulk RNA-seq benchmark

The real bulk benchmark used the raw_gene_counts.tsv retinal count matrix distributed with CoTRA. The dataset contained 55,291 genes and 12 samples comprising four WT and eight rd10 retinas. The rd10-versus-WT comparison was evaluated independently using the DESeq2 and edgeR implementations available through CoTRA.

DESeq2 retained 19,982 genes for statistical testing and identified 2,518 genes meeting the applied adjusted P-value and absolute log2 fold-change thresholds. Median analytical runtime was 12.93 s (Q1–Q3: 12.78–12.96 s), and median peak-memory consumption was 0.88 GB. The differential expression step required approximately 8.25 s.

edgeR retained 18,922 genes and identified 2,322 significant genes. Median analytical runtime was 2.53 s (Q1–Q3: 2.52–2.53 s), median peak-memory consumption was 0.65 GB, and the differential-expression step required approximately 1.92 s.

Differences in genes retained or identified reflect method-specific filtering and statistical procedures and were not interpreted as evidence that either method provides superior statistical performance. Complete runtime, memory, run-variability, genes-tested, and significant-gene statistics are provided in **S4 Table**, with step-level timings in **S7 Table.**

**Synthetic scRNA-seq benchmark**

Synthetic sparse scRNA-seq datasets contain 20,000 genes and 2,500, 5,000, 10,000, 25,000, or 50,000 cells. The simulation contained 12 predefined populations with 150 designated marker genes per population, corresponding to 1,800 population-associated markers. Fixed nested subsets were generated from the same underlying simulation so that increasing dataset sizes represented progressively larger subsets rather than independently generated datasets. Complete simulation parameters and generation scripts are provided in **S1 Code**.

The timed core workflow comprised Seurat object construction and quality-control metric calculation, LogNormalize normalization using a scale factor of 10,000, selection of 2,000 highly variable genes using the vst method, scaling, PCA, UMAP, nearest-neighbor graph construction, and Louvain clustering. Fifty principal components were calculated, and PCs 1-30 were used for downstream analysis. Neighbor graphs were generated using k.param = 20, and clustering used Louvain algorithm 1, resolution 0.5, and random seed 1234.

Because this experiment assessed computational scaling rather than clustering accuracy, marker-analysis timing was evaluated separately using the 12 known simulated population labels. FindAllMarkers() was run using the Wilcoxon test with only.pos = TRUE, min.pct = 0.10, a log2 fold-change threshold of 0.25, and a maximum of 500 cells per population. The reported combined value therefore represents the core-workflow plus truth-label marker-analysis workload, rather than a cluster-derived end-to-end marker result.

Median core-workflow runtimes were 12.36, 23.16, 42.44, 103.39, and 291.05 s for 2,500, 5,000, 10,000, 25,000, and 50,000 cells, respectively. Median marker-analysis times were 0.91, 1.36, 2.04, 4.22, and 7.78 s, resulting in combined workloads of 13.29, 24.54, 44.49, 107.62, and 299.35 s.

Median peak-memory consumption increased from 0.67 GB at 2,500 cells to 0.97, 1.40, 2.75, and 5.10 GB at 5,000, 10,000, 25,000, and 50,000 cells, respectively. All 25 measured runs completed successfully.

At 50,000 cells, PCA was the largest computational component, requiring approximately 179.5 s, followed by Louvain clustering (~53.2 s), UMAP (~37.4 s), and nearest-neighbor graph construction (~12.2 s). Complete core-workflow, marker-analysis, combined-runtime, peak-memory, cluster-count, and marker statistics are reported in **S5 Table**, with individual step-level timings in **S7 Table**. Run-level outputs are provided in **S1 Data**, processed benchmark summaries and figure-source data in **S2 Data**, and synthetic data generation and benchmark scripts in **S1 Code**.

#### Real retinal scRNA-seq benchmark

The real scRNA-seq performance benchmark used the WT non-treated and vehicle-treated rd10 samples from GSE234797; the TMB-treated rd10 sample was excluded. Processed GSE234797 cell identifiers and metadata were used to identify the cells represented in the processed dataset and to map those identifiers back to the corresponding original 10x Genomics HDF5 matrices, thereby recovering their raw integer counts.

This procedure recovered 4,478 WT and 5,952 vehicle-treated rd10 cells, corresponding to 10,430 cells in total. Of the 18,545 genes represented in the processed gene list, 18,533 were present in both raw HDF5 matrices and were retained for benchmarking. Sparse matrix representation was maintained throughout the scRNA-seq workflow.

Condition-stratified nested subsets containing 2,500, 5,000, 7,500, and 10,000 cells were generated in addition to the complete 10,430-cell dataset while approximately preserving the WT-to-rd10 composition. Fifty principal components were calculated, and PCs 1-7 were used for UMAP, nearest-neighbor graph construction, and clustering, consistent with the principal-component selection used for the retinal CoTRA analysis. Marker analysis was performed using the graph-based clusters inferred independently from each benchmark dataset.

Median combined core-workflow and marker-analysis runtimes were 15.72, 26.01, 36.29, and 47.29 s for the 2,500-, 5,000-, 7,500-, and 10,000 cell datasets, respectively. Median peak-memory consumption increased from 1.21 GB to 2,500 cells to 1.96, 2.65, and 3.37 GB for the corresponding larger subsets.

For the complete 10,430 cell dataset, median core-workflow runtime was 25.87 s, and median marker-analysis runtime was 20.89 s, resulting in a combined median runtime of 46.68 s and median peak-memory consumption of 3.36 GB. The benchmark produced 17 graph-based clusters. Cluster number was recorded descriptively rather than treated as a computational-performance criterion because graph-based clustering can vary with software version, input representation, and analytical settings.

The slightly shorter combined runtime for the complete 10,430-cell dataset compared with the 10,000-cell subset arose primarily from dataset-size-dependent behavior of the underlying UMAP implementation. Package defaults were retained to represent the current CoTRA execution environment rather than modifying UMAP parameters solely to force monotonic runtime scaling.

All 25 measured real scRNA-seq runs completed successfully, with total-runtime coefficients of variation below 1% for every tested dataset size. Complete cell composition, core-workflow, marker-analysis, combined-runtime, memory, cluster-count, and marker statistics are reported in **S6 Table**, with step-level timings in **S7 Table**. Run-level outputs are provided in **S1 Data**, processed benchmark summaries in **S2 Data**, and scripts for processed-cell reconstruction, benchmark execution, and summarization in **S1 Code**.

Overall, these experiments characterize the runtime, memory requirements, scalability, and execution stability of CoTRA and are distinct from the implementation-concordance analyses described below.

### SUPPLEMENTARY IMPLEMENTATION VALIDATION

#### Validation strategy and concordance metrics

Implementation validation assessed whether CoTRA reproduces the corresponding direct scripted R-package calculations when identical input data and analytical settings are used. This analysis therefore evaluates **implementation fidelity**, rather than attempting to revalidate the statistical methodology of DESeq2, edgeR, or Seurat.

For result sets A and B, overlap was quantified using the Jaccard index,

$$J(A,B) = \frac{|A \cap B|}{|A \cup B|}$$

where J=1 indicates identical sets.

For continuous quantities such as log2 fold-change estimates and PCA scores, Pearson and, where appropriate, Spearman correlations were calculated.

For scRNA-seq cluster assignments, agreement was quantified using the adjusted Rand index (ARI),

$$\mathrm{ARI} = \frac{\sum_{ij}\binom{n_{ij}}{2} - \frac{\left(\sum_i \binom{a_i}{2}\right)\left(\sum_j \binom{b_j}{2}\right)}{\binom{n}{2}}}{\frac{1}{2}\left[\sum_i \binom{a_i}{2} + \sum_j \binom{b_j}{2}\right] - \frac{\left(\sum_i \binom{a_i}{2}\right)\left(\sum_j \binom{b_j}{2}\right)}{\binom{n}{2}}}$$

where $n_{ij}$ represents the number of cells shared by cluster i in one partition and cluster j in the other, while $a_i$ and $b_j$ are the corresponding marginal cluster sizes.

Normalized mutual information (NMI) was calculated as

$$\mathrm{NMI}(C,D) = \frac{I(C;D)}{\sqrt{H(C)\,H(D)}}$$

where I(C;D) is the mutual information between cluster assignments C and D, and H denotes entropy. ARI and NMI values of 1 indicate identical cell partitions.

Scripts used for the implementation-concordance analyses are provided in **S2 Code** and under validation/Implementation_validation_code/. Machine-readable outputs underlying **S8 and S9 Tables** are provided in **S4 Data** and under validation/Implementation_validation_outputs/.

**Direct CoTRA versus scripted DESeq2**

Bulk implementation concordance was evaluated using the raw_gene_counts.tsv retinal test dataset. The same 55,291 gene count matrix, WT/rd10 sample assignments, filtering procedures, rd10-versus-WT contrast, and significance thresholds were used in CoTRA and in a direct scripted DESeq2 analysis.

Both implementations retained 19,982 genes and identified exactly 2,518 significant genes. Log2 fold-change estimates were identical, the significant-gene Jaccard index was 1.000, Pearson and Spearman correlations for log2 fold-change estimates were 1.000, and direction of change was concordant for 100% of significant genes.

The raw direct DESeq2 output contained 127 genes for which DESeq2 returned undefined (NA) raw and adjusted P values. Consistent with the current CoTRA implementation, these values were assigned a value of 1 before significance filtering to permit stable downstream classification and visualization. This post-processing did not change the significant-gene set. After applying the documented CoTRA post-processing step, the maximum absolute numerical difference between compared exported values was zero.

Complete DESeq2 implementation-concordance statistics are included in **S8 Table**, with machine-readable run-level comparisons in **S4 Data**.

**Direct CoTRA versus scripted edgeR**

The same bulk RNA-seq dataset, sample grouping, contrast, filtering procedure, and significance thresholds were used for the edgeR comparison.

Both implementations retained 18,922 genes and identified exactly 2,322 significant genes. The logFC, logCPM, quasi-likelihood F statistic, raw P-value, and FDR values were numerically identical for all retained genes. The significant-gene Jaccard index was

1.000, log2 fold-change Pearson and Spearman correlations were 1.000, and directional concordance was 100%.

The five independently executed scripted analyses produced the same analytical results in every run. Complete bulk implementation validation statistics are summarized in **S8 Table**, and the corresponding machine-readable outputs are provided in **S4 Data**.

**Direct CoTRA versus scripted Seurat**

Single-cell implementation validation used the two 10x Genomics HDF5 files distributed with the CoTRA repository:

- GSM7474906_Wild_Type_non_treated_feature_bc_matrix.h5
- GSM7474907_Rd10_Female_vehicle_feature_bc_matrix.h5.

The matrices contained 32,285 genes and, before additional user-defined filtering, 7,217 WT and 7,889 rd10 cells, corresponding to 15,106 cells in total.

For implementation validation, all cells contained in the supplied Cell Ranger-filtered HDF5 matrices were retained so that the comparison tested the direct relationship.

Identical input + Identical parameters ⟶ CoTRA versus direct Seurat

Both workflows used LogNormalize normalization with a scale factor of 10,000, selection of 2,000 highly variable features using vst, scaling of the variable genes, computation of 50 principal components, PCs 1-7 for downstream graph construction, FindNeighbors() with k.param = 20, and Louvain clustering using algorithm 1, resolution 0.5, and random seed 1234. Marker analysis used the Wilcoxon test with only.pos = TRUE, min.pct = 0.10, a log2 fold-change threshold of 0.25, and max.cells.per.ident = 500.

CoTRA and direct Seurat selected the same 2,000 highly variable genes, corresponding to an HVG Jaccard index of 1.000. PCA score correlations for PCs 1-7 were 1.000 for every component.

Both workflows assigned all 15,106 cells to the same 16 graph-based clusters. Cell-level cluster agreement therefore yielded ARI = 1.000 and NMI = 1.000. Cluster-label mapping and the complete cell-level contingency matrix are provided in **S4 Data**.

Marker analysis generated 38,491 gene–cluster pairs in both workflows. The complete marker-set Jaccard index was 1.000, the significant marker-set Jaccard index was 1.000, and average marker log2 fold-change values had Pearson and Spearman correlations of 1.000.

Complete scRNA-seq implementation-concordance results are summarized in **S9 Table**. PCA concordance values, cluster contingency matrices, cluster-label mappings, and machine-readable scRNA-seq validation results are provided **in S4 Data**, while the direct Seurat validation scripts are included in **S2 Code**.

Together, these analyses demonstrate that the CoTRA DESeq2, edgeR, and Seurat workflows reproduce their corresponding direct scripted implementations when identical inputs and analytical settings are used.

### SUPPLEMENTARY TEST-DATA AND REPRODUCTION

#### Bulk RNA-seq test dataset

The primary bulk RNA-seq test dataset distributed with CoTRA is raw_gene_counts.tsv. The matrix contains 55,291 genes and 12 retinal samples comprising four WT and eight rd10 samples. The dataset is intended to allow users to test count-matrix import, sample grouping, exploratory analysis, differential-expression analysis, annotation, functional enrichment, visualization, and export functions.

Under the computational environment reported in **S2 Table** and the fixed validation settings summarized in **S10 Table**, the expected differential-expression outputs are 19,982 genes tested and 2,518 significant genes for DESeq2 and 18,922 genes tested and 2,322 significant genes for edgeR. These values are provided as reproducibility checks for the specified configuration rather than immutable outputs, because changing filtering, significance thresholds, package versions, or other user-configurable parameters can alter the results.

The dataset characteristics, recommended test settings, and expected outputs are summarized in **S10 Table**.

#### scRNA-seq test datasets

Two 10x Genomics HDF5 matrices are distributed for scRNA-seq testing:

- GSM7474906_Wild_Type_non_treated_feature_bc_matrix.h5
- GSM7474907_Rd10_Female_vehicle_feature_bc_matrix.h5.

The WT matrix contains 32,285 genes and 7,217 cells, while the rd10 matrix contains 32,285 genes and 7,889 cells. Together, the files provide a realistic two-sample scRNA-seq input containing 15,106 cells.

The files can be used to test multiple-H5 import, biological sample naming, experimental-condition assignment, barcode prefixing, object construction, quality-control metric calculation, normalization, highly variable feature selection, PCA, UMAP, graph construction, clustering, marker detection, cell-type annotation, and subsequent single-cell analysis modules.

For the fixed implementation-validation configuration, no additional user-defined QC filtering was applied. LogNormalize with a scale factor of 10,000, 2,000 vst highly variable genes, 50 principal components, PCs 1-7, k.param = 20, Louvain algorithm 1, resolution 0.5, and random seed 1234 produced 16 clusters. Marker analysis using the Wilcoxon test, only.pos = TRUE, min.pct = 0.10, log2FC threshold 0.25, and max.cells.per.ident = 500 produced 38,491 marker gene–cluster pairs. These values represent expected outputs for the specified validation environment and settings.

The dataset characteristics and expected validation outputs are summarized in **S10 Table**.

**Reproduction instructions**

For the bulk test, users should import raw_gene_counts.tsv, define WT as the reference group and rd10 as the comparison group, and run DESeq2 or edgeR using the settings specified in **S10 Table**. The resulting complete differential-expression tables can be compared with the reference outputs in **S4 Data**.

For the scRNA-seq test, the two HDF5 files should be imported as separate biological samples, assigned WT and rd10 sample/condition labels, and combined using unique sample-prefixed cell barcodes. To reproduce the implementation-validation experiment, all cells contained in the supplied filtered HDF5 matrices should be retained, and the fixed Seurat parameters listed above should be applied. Resulting HVGs, PCA coordinates, cluster assignments, and marker tables can then be compared with the corresponding reference outputs in **S4 Data**.

Scripts for reproducing the direct implementation comparisons are available in **S2 Code** and in the GitHub directories validation/Implementation_validation_code/bulk/ and validation/Implementation_validation_code/scRNA/. Reference validation outputs are available under validation/Implementation_validation_outputs/.

The complete computational-performance resources are separately organized under benchmarking/code/ and benchmarking/outputs/, allowing users to reproduce the computational analyses underlying **Fig 6** and **S2-S7 Tables** without mixing benchmarking resources with implementation-validation outputs.

### SUPPLEMENTARY SOFTWARE AVAILABILITY, INSTALLATION AND REPRODUCIBILTY RESOURCES

CoTRA methods and configurable settings are summarized in **S1 Table**, while the computational environment, benchmark outputs, validation resources, and reproducibility test settings are provided in **S2-S10 Tables** and **S1-S4 Data/S1-S2 Code**. The package requires R ≥ 4.4.0. Installation instructions are provided in the repository for Linux/Ubuntu, Windows, and macOS, together with platform-specific dependency guidance.

CoTRA can be installed directly from GitHub using:

```
remotes::install_github("UmairSeemab/CoTRA", dependencies = TRUE)
```

Additional analysis dependencies can be installed where required using:

```
CoTRA::install_cotra_dependencies()
```

The graphical interface is launched with:

```
CoTRA::runCoTRA()
```

During execution, users can select an output directory for generated results. When no output directory is explicitly selected, CoTRA uses ~/CoTRA_Results as the default location. Depending on the analysis module, generated outputs can include tables, figures, reports, gene lists, ZIP archives, processed RDS or Seurat objects, CoTRA project/session files, and session information.

Representative real datasets used in the present study are available as reproducible test inputs. The bulk RNA-seq test dataset is provided as data/bulkRNA/raw_gene_counts.tsv. The scRNA-seq test inputs are the 10x Genomics HDF5 files data/scRNA/GSM7474906_Wild_Type_non_treated_feature_bc_matrix.h5 and data/scRNA/GSM7474907_Rd10_Female_vehicle_feature_bc_matrix.h5. The bulk dataset supports testing of import, sample grouping, exploratory analysis, differential-expression analysis, annotation, enrichment, visualization, and export. The two HDF5 files support testing of multi-sample import, sample and condition assignment, quality control, normalization, highly variable feature selection, dimensionality reduction, clustering, marker analysis, annotation, and downstream single-cell analysis.

The parameter settings used to generate the results reported in this study and the corresponding expected outputs are provided in the Supplementary tables. These reference outputs are intended to facilitate installation checks and reproducibility testing; exact downstream results may change when users modify configurable parameters or use different versions of the underlying analytical packages.

Computational-performance resources are organized in the repository under benchmarking. The benchmarking/code/ directory contains the scripts used for synthetic-data generation, real-data preparation, benchmark execution, result summarization, and figure generation, while benchmarking/outputs/ contains machine-readable benchmark summaries, run-level measurements, step-level timings, and resource-usage files underlying the reported performance results.

Implementation-concordance resources are organized separately under validation. The validation/Implementation_validation_code/ directory contains the scripts used to compare CoTRA with direct scripted DESeq2, edgeR, and Seurat analyses, and validation/Implementation_validation_outputs/ contains the corresponding machine-readable concordance results underlying the implementation-validation tables. These resources allow the reported agreement statistics to be independently inspected and reproduced.

CoTRA currently begins with processed expression data and does not perform FASTQ preprocessing, read alignment, or read quantification. Bulk and single-cell RNA-seq are implemented as separate analytical branches within the same application and are not currently combined through a joint cross-modality workflow. Large scRNA-seq analyses remain subject to the computational and memory requirements of the underlying R and Seurat implementations. In addition, statistical interpretation of differential expression, differential abundance, pathway activity, and related analyses depends on the experimental design, biological replication, and assumptions of the selected underlying methods.

The public GitHub issue tracker provides a mechanism for reporting software problems, requesting features, and proposing contributions. The modular R/Shiny architecture is intended to facilitate addition of further analytical methods and reproducibility features as transcriptomic-analysis practices evolve.

### SUPPORTING INFORMATION

**S1 Fig. CoTRA graphical user interface.** Representative CoTRA interface showing workflow navigation, user-configurable analytical parameters, visualization of results, download options, and integrated interpretation guidance.

**S2 Fig. Feature comparison of CoTRA with representative graphical RNA-seq analysis platforms.** Heatmap comparing CoTRA and 14 other platforms across 49 predefined criteria covering deployment, usability, input handling, bulk and single-cell RNA-seq workflows, visualization, biological interpretation, advanced analyses, export, reporting, and reproducibility.

**S3 Fig. Quality-control characteristics of WT and rd10 mouse retinal single-cell RNA-seq data.** Violin plots showing distributions of detected genes per cell, total RNA counts, mitochondrial transcript percentage, and ribosomal transcript percentage.

**S4 Fig. Cluster-associated marker gene expression in the CoTRA analysis.** Dot plot showing the three highest-ranking marker genes for each of the 16 identified clusters; dot size represents the proportion of expressing cells and color represents average expression.

**S5 Fig. UMAP visualization of representative cluster-associated genes.** Feature plots showing expression distributions of representative retinal marker genes across the UMAP embedding.

**S1 Table. CoTRA analysis methods, default implementations, study settings, and user-configurable parameters.**

**S2 Table. Computational environment used for CoTRA benchmarking.**

**S3 Table. Synthetic bulk RNA-seq computational benchmark results.**

**S4 Table. Real retinal bulk RNA-seq computational benchmark results.**

**S5 Table. Synthetic scRNA-seq computational benchmark results.**

**S6 Table. Real retinal scRNA-seq computational benchmark results.**

**S7 Table. Step-level computational timing results.**

**S8 Table. Concordance between CoTRA and direct scripted bulk differential-expression implementations.** Concordance results for DESeq2 and edgeR analyses performed using identical input data and analytical settings.

**S9 Table. Concordance between CoTRA and direct scripted Seurat analysis.** Concordance of highly variable genes, PCA coordinates, cluster assignments, and marker-analysis results under matched input data and parameters.

**S10 Table. CoTRA test datasets, fixed validation settings, and expected outputs.** Characteristics of the supplied bulk and scRNA-seq test datasets together with the fixed settings and expected outputs used for reproducibility checks.

**S11 Table. Feature comparison of CoTRA and 14 representative transcriptomic analysis platforms across 49 predefined criteria.** Includes the criterion-level comparison, definitions, evidence sources, software versions or releases, and evaluation information.

**S1 Data. Run-level measured computational-benchmark outputs.** Run-level runtime, step-level timing, and GNU time -v resource measurements underlying the reported computational-performance results.

**S2 Data. Processed computational-benchmark summaries and figure-source data.** Machine-readable benchmark summaries, computational-environment information, step-level summaries, and source data used to generate the computational-performance figure.

**S3 Data. Preliminary computational-benchmark viability runs.** Preliminary runs performed to confirm successful execution before the five measured replicates; these runs were excluded from all reported benchmark statistics.

**S4 Data. Machine-readable implementation-validation outputs.** Bulk and scRNA-seq concordance summaries, run-level comparisons, PCA concordance results, cluster contingency matrices, and cluster-label mappings underlying S8 and S9 Tables.

**S1 Code. CoTRA computational benchmarking scripts.** Scripts used for synthetic-data generation, real-data preparation, computational benchmark execution, result summarization, and computational-performance figure generation.

**S2 Code. CoTRA implementation-validation scripts.** Scripts used to compare CoTRA with direct DESeq2, edgeR, and Seurat implementations using matched input data and analytical settings.
